# Funding deadlines act as a zeitgeber for science

**DOI:** 10.64898/2026.10.10.758253

**Authors:** Haining Wang, Jing Su

## Abstract

Funders decide which science is paid for. Whether they also shape when it becomes public is hard to test. Across 376,542 preprints on bioRxiv and medRxiv, the US share rose by 2.3 to 2.7 percentage points in the four weeks before due dates for the main research grant of the National Institutes of Health (NIH) and dropped on the date. About one in ten US preprints in those weeks was attributable to the deadline, and fields that depend least on NIH rose least. Among 15,906 funded investigators, the share of preprints posted in the four weeks before their own deadline was 9% higher than in other four-month cycles, and these preprints resembled the later grant without being more highly cited. Repeated before every application due date, this individual surge matches the national peak’s timing within two weeks. NIH’s application calendar, like its funding, shapes when US biomedical science becomes public.

## Introduction

When knowledge becomes public is not a private decision. Merton described scientific disclosure as governed by institutions that reward being first (1), and the economics of science treats the timing of disclosure as a response to the rewards attached to it (2, 3). These accounts have been difficult to test because the moment a result became public was usually recorded only as the date of journal publication, which is slow and variable. Preprints changed that. A first version posted to bioRxiv or medRxiv carries a date chosen by its authors, and a growing share of biomedical research now reaches readers first in this form (4, 5). We ask what decides when a first version appears.

A funder is a candidate. The National Institutes of Health (NIH), the largest public funder of biomedical research, receives most investigator-initiated applications on fixed dates three times a year: new applications for the R01, its main research grant, on 5 February, 5 June and 5 October. Career awards (K), smaller and exploratory grants (R03 and R21), trainee fellowships (F), resubmissions and, until recently, AIDS-designated applications have their own dates. Reviewers weigh the scientific background of a proposal, including preliminary data, and its feasibility, criteria that remain under the 2025 simplified framework (6). Since the 25 May 2017 due date, applicants have been able to cite preprints if the citation carries the preprint’s digital object identifier (7). bioRxiv and medRxiv issue the identifier once the author approves posting, and it resolves only when the screened preprint is public (8). A result meant to count in an application must therefore be public before the due date.

We frame the consequence as entrainment. In chronobiology a zeitgeber is an external cue that sets the phase of a rhythm without supplying the rhythm itself (9). In organizations, external cycles such as budget calendars set the pace of activity in the groups that depend on them (10, 11), and people reorganize their work in anticipation of how it will be judged (12). A funder’s calendar should entrain disclosure when three conditions hold together: a clock (fixed receipt dates), a currency (reviewers who value recent results) and an exchange rule (a way for public results to count in the application). By entrainment we mean the recurrent alignment of disclosure with an external evaluation calendar. The rhythm the calendar sets in phase is each investigator’s own slow cycle of output around grants, *h*_0_*_i_*(*t*) in the model below. A minimal model makes the conditions explicit. Investigator *i* posts first versions on day *t* at rate

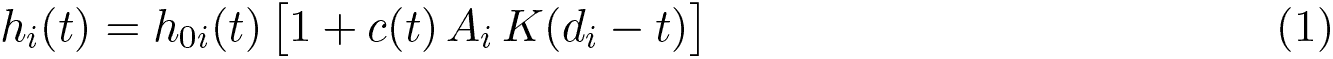

where *h*_0_*_i_*(*t*) is the investigator’s own slow cycle of output, *d_i_* is the investigator’s next due date, *K*(*u*) is the clock, a short bump (kernel) over the days *u* before a due date that is positive for a few weeks and zero once the date has passed, *A_i_ ≥* 0 is the investigator’s coupling to the funder (the currency), and *c*(*t*), the exchange rule, is 1 when a public result can count in an application and 0 otherwise. For a field *f* whose US authors depend on the funder with weight *D_f_* and apply on due dates spread with density *p_d_*, the change in the US share *S_f_* at cycle phase *ϕ*, the number of days to the next R01 due date, is

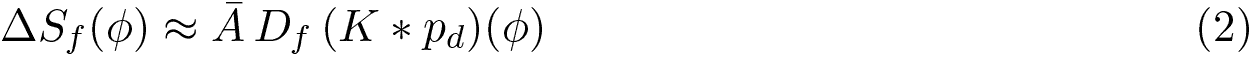

where *Ā* is the average coupling multiplied by *S_f_* (1 *− S_f_*) and *K ∗ p_d_* is the kernel convolved with the spread of due dates, that is, the bump summed over all due dates (derivation in text S2). The model predicts (i) a drop on the due date, because *K* ends there, (ii) an excess confined to the weeks before an investigator’s own due date, (iii) a national (aggregate) profile given by the individual kernel placed before every due date and (iv) a rise that grows with dependence on the funder. The data identify *K* and its placement on the calendar.

NIH’s four-month cycle repeats three times a year, so a rhythm with that period could also come from the academic calendar, holidays or conferences. The predictions separate the funder’s clock from these. A season changes smoothly across a date and cannot drop on it, and an investigator’s own due date moves between cycles while the academic calendar does not. We linked 171,167 NIH awards to 67,184 principal investigators and their preprints. US preprinting dropped on the R01 due date, individual investigators posted the science of their applications in the two weeks before their own due date, that individual excess placed the national peak within a week of where it is observed on bioRxiv and within two weeks on medRxiv, and fields that depend least on NIH rose least.

## Results

### US preprinting rises before NIH due dates and drops on them

If NIH’s calendar sets when science goes public, the share of preprints from US authors should rise before its due dates and drop on the date NIH uses. US preprinting does both.

We placed each day on the NIH cycle by its distance to the next R01 due date and compared the window from 28 days before through the due date (the R01 window) with the rest of the four-month cycle, with standard errors clustered by cycle (30 bioRxiv and 22 medRxiv cycles). In this window the US share of first versions rose by 2.26 percentage points on bioRxiv (SE 0.33; mean US share 40.7%) and by 2.70 points on medRxiv (SE 0.53; mean 34.5%) (table S3), most of it in the final three weeks (table S4 and fig. S1A). The rise came mostly from more US work. On the log scale, where 0.1 is roughly a 10% increase, daily US first versions rose by 0.150 on bioRxiv (SE 0.018) and by 0.170 on medRxiv (SE 0.024), against 0.056 (SE 0.016) and 0.053 (SE 0.019) for non-US first versions (table S5). Taking the non-US change as the baseline, about one in eleven US first versions posted in the window on bioRxiv, and one in nine on medRxiv, is there because of the deadline. The rise is visible in the raw counts of most years (Fig. 1A) and in the window of each of the three annual cycles (table S6). It was clearest from 2021. From 2017 to 2020 the yearly estimates were smaller and less precise, and one was negative (table S1); before 2017 no rhythm was detectable, in a comparison too small to detect a rise of this size (fig. S2).

**Fig. 1.**
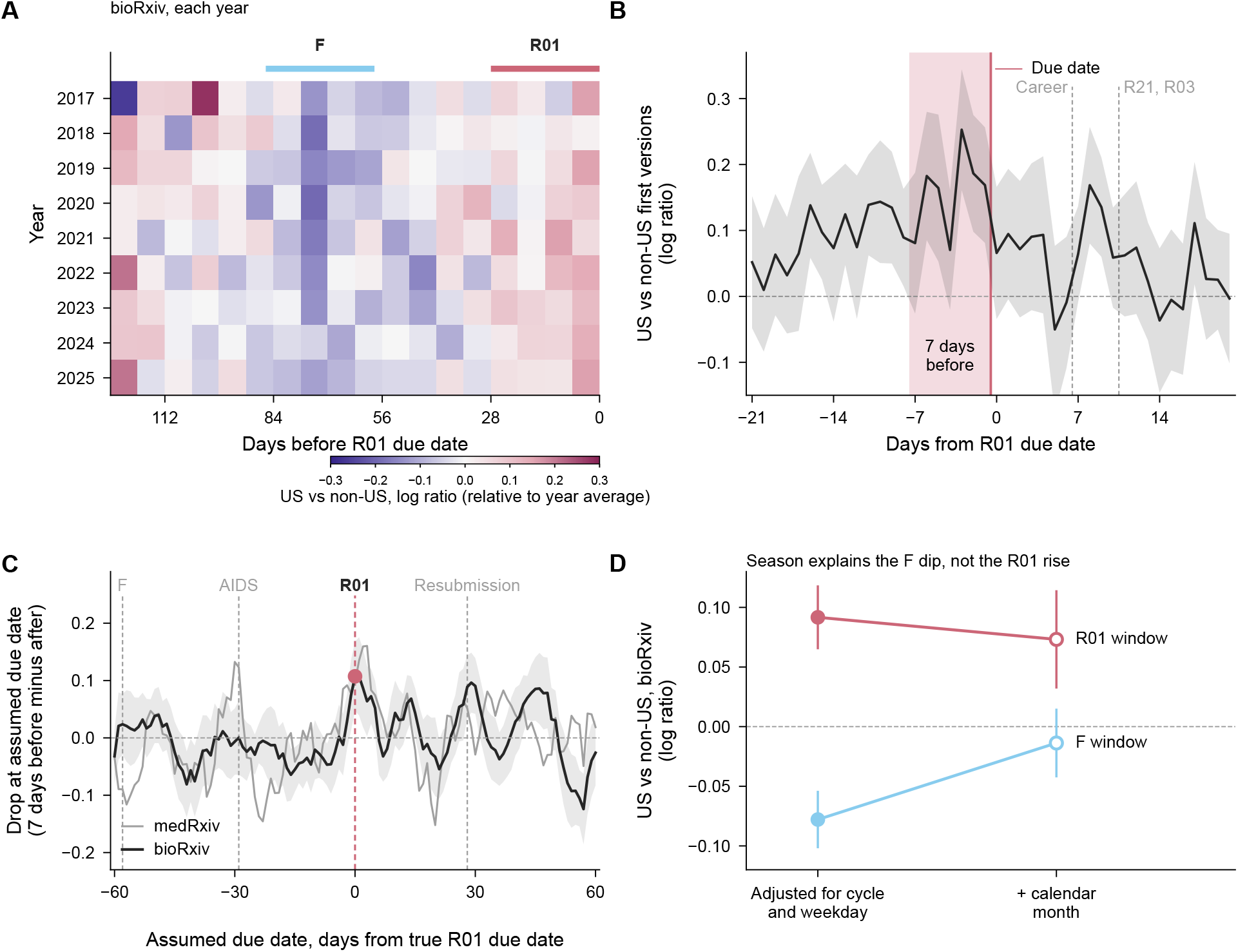
US preprinting rises before NIH due dates and drops on them. (**A**) Raw data. For each year and each 7-day bin of the NIH cycle, the log ratio of US to non-US first versions on bioRxiv, relative to that year’s average (centered within the year); rose cells, more US preprints. The bars above mark the 28 days before R01 (rose) and fellowship (F; light blue) due dates. The rise before R01 due dates recurs in most years (table S1). (**B**) Day by day around the R01 due date (vertical line), the log ratio of US to non-US daily first versions on bioRxiv relative to other days (Poisson model with day fixed effects, which compares US with non-US preprints on the same day). Shading marks the 7 days before the due date; ticks mark the due dates of career awards and of R21 and R03 grants. The ratio peaks just before the due date and drops on it. A second rise 8 to 9 days after the due date falls between the due dates of career awards and of R21 and R03 grants (text S1). (**C**) The drop, the 7 days before an assumed due date minus the 7 days from it, as the assumed date moves from 60 days before to 60 days after the true R01 due dates (bioRxiv, black, with 95% confidence band; medRxiv, gray). The drop is largest at the true dates (rose); ticks mark other NIH due dates. Rank tests against shifted dates are in table S2. (**D**) Rise of US over non-US daily first versions on bioRxiv in the R01 and F windows (window coefficients), adjusted for cycle and weekday (fixed effects; filled) and also for calendar month (open). Adjusting for calendar month removes the F-window dip and leaves most of the R01-window rise. Error bars and bands, 95% confidence intervals.

Day by day, the ratio of US to non-US first versions was highest one to three days before the R01 due date (0.17 to 0.25 on the log scale) and fell to 0.07 on the due date (Fig. 1B). Because a cited preprint must be public before the application is due, any posting the deadline pulls forward should stop on the due date. The drop between the 7 days before and the 7 days from the due date was 0.107 on bioRxiv (SE 0.031) and 0.096 on medRxiv (SE 0.053). With every due date moved by 14 to 60 days in either direction, the drops averaged zero, and none of the 94 shifted dates gave a drop as large as the true one on bioRxiv (rank *P* = 0.011; medRxiv, 0.032) (Fig. 1C and table S2). The drop also follows the date NIH actually uses. When 5 February, 5 June or 5 October fell on a weekend, NIH accepted applications on the following Monday. The weekend days in between behaved like the days before a deadline and not like the days after it. Their estimates were 0.133 on bioRxiv (SE 0.103) and 0.421 on medRxiv (SE 0.101), against 0.139 and 0.116 for the 7 days before the published date and 0.027 and *−*0.001 for the days after it (table S7). The drop was largest in February, the cycle with no federal holiday near the due date (0.172, against 0.100 in June and 0.057 in October). The drop held with a linear trend on each side of the date and without East Asian authors or holiday weeks in the comparison (text S1).

Fixed effects for each group (US or non-US) by calendar month leave only differences within a month and so show what the season contributes. With them, the R01-window coefficient on bioRxiv daily counts was 0.073 (SE 0.021), against 0.092 (SE 0.014) without them, whereas a dip in the fellowship window (the 28 days before F due dates) on the opposite side of the cycle disappeared (*−*0.014, SE 0.015, against *−*0.078, SE 0.012) (Fig. 1D and table S8). The fellowship-window dip is therefore seasonal. The June window, which ends with the US academic year, carries an annual component of the same kind: its window effect is the largest of the three cycles, while its drop at the due date is not (tables S6 and S2).

### Investigators post in the two weeks before their own due date, whichever date it is

Within a person, the academic calendar is the same in every cycle, but the investigator’s own due date is not (Fig. 2A). An investigator who posts to meet a deadline should post more in the weeks before the date they face, and the excess should move when that date moves.

**Fig. 2.**
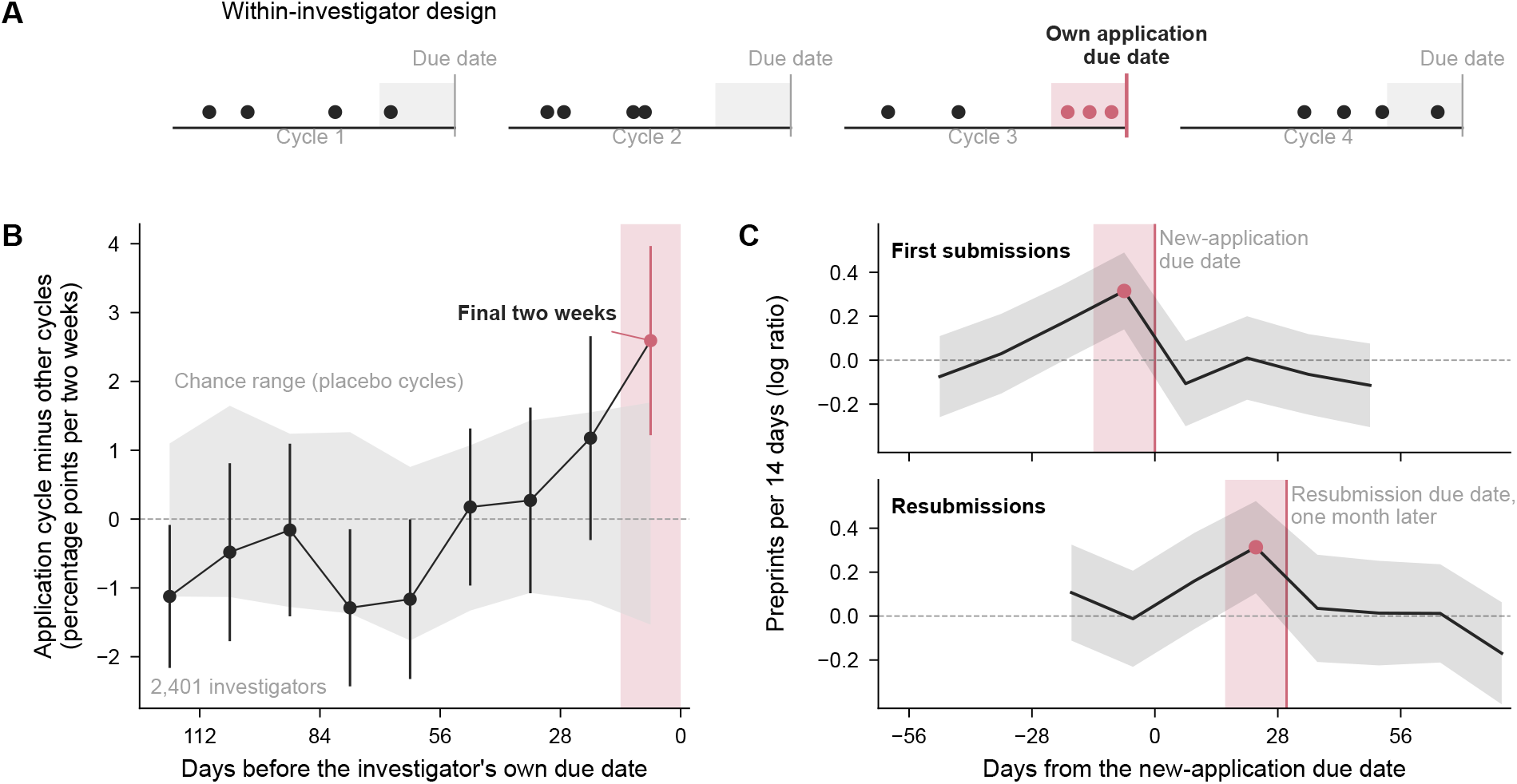
Investigators post in the two weeks before their own due date, whichever date it is. (**A**) The within-investigator design. Each cycle has a due date and the same academic calendar, and in one cycle the investigator submits an application. The comparison is the share of the investigator’s preprints posted in the 28 days before the due date (shaded) in that cycle against the same investigator’s other cycles. (**B**) Difference in the share of a cycle’s preprints posted in each two-week period before the due date, application cycles minus other cycles (2,401 investigators). Gray band, the range expected by chance: 2.5th to 97.5th percentiles when each application cycle is replaced by another of the investigator’s cycles (placebo cycles, 200 draws). Only the final two weeks leave the band. The differences sum to zero over the cycle, so earlier periods sit slightly below zero. (**C**) Preprint counts of linked investigators in 14-day bins around their own due date, compared within each application and within calendar time (event and calendar fixed effects): funded first submissions around the new-application due date (top) and funded resubmissions around the resubmission due date (bottom), shifted 30 days later, the average gap between the two dates. Both rise in the two weeks before their own date (rose) and not after it. Bands and error bars, 95% confidence intervals.

From NIH RePORTER, NIH’s public database of awards, we inferred the due date of each funded application from its award notice month and grant type (activity code), and linked principal investigators to their preprints only through papers that acknowledged the investigator’s own award. For each investigator and each four-month cycle we computed the share of the cycle’s last-author preprints posted in the 28 days before the due date, and compared cycles in which the investigator submitted a funded new application with the same investigator’s other cycles, with fixed effects for investigator and calendar cycle. An investigator may choose to apply in the cycle in which results are ready. What we estimate is therefore the timing of posting within the cycle the investigator chose, among funded, linked investigators who posted preprints. Readiness that is independent of the deadline predicts posting at any time before submission, and it does not predict an excess confined to the final weeks.

Among 15,906 investigators, the share was 2.3 percentage points higher in the application cycle (SE 1.0), a relative increase of 9% (log ratio 0.087, SE 0.039), resting on about 1,000 pre-deadline preprints. Placebo cycles drawn among each investigator’s other cycles centered on zero, and the observed estimate exceeded 98% of them (*P* = 0.015). Linking investigators to authors only through works other than preprints and their journal versions gave the same estimate (0.087, SE 0.039), and so did comparing each application cycle only with the investigator’s other cycles in the same months of the year (0.090, SE 0.047), so the estimate does not depend on links made through the outcome or on a season peculiar to the investigator (tables S9 and S10). For the 2,401 investigators with preprints in both kinds of cycle, the share posted in the last two weeks was 2.6 points higher in the application cycle (SE 0.7), about one-fifth above a base of 12.8% and outside the range of placebo cycles, whereas every earlier two-week period stayed within that range (Fig. 2B).

The excess follows the date the investigator faces. NIH receives resubmissions about a month after new applications (5 March, 5 July and 5 November for R01 grants), and a resubmission may add new data in reply to the first review. In counts of each investigator’s preprints, funded first submissions rose by 0.315 on the log scale (SE 0.089) in the 14 days before the new-application due date, and funded resubmissions rose by 0.313 (SE 0.107) in the 14 days before the resubmission due date a month later (Fig. 2C and table S11). Neither rose detectably after its date (*−*0.107, SE 0.099, and 0.035, SE 0.124), so first submitters showed no sign of posting the work of their applications after submitting them.

A due date inferred in the wrong cycle and unfunded applications hidden among the reference cycles both pull the estimate toward zero (text S3 and table S12). Because only funded applications are observed, posting that raised the chance of funding would inflate it, so the net bias has no known sign. Selection of this kind would not confine the excess to the final two weeks, however. A preprint posted two months before a due date is as citable as one posted in the final two weeks, so selection on posting would have to favor only the latter.

### One surge, repeated on every due date, places the national peak

If applicants set the national rhythm, their individual two-week surge (the excess in Fig. 2B), placed before every NIH new-application due date, should put the national peak where it is observed, and fields whose researchers depend least on NIH should rise least.

NIH’s new-application due dates are spread over three weeks of each cycle (5th for R01, 12th for career awards, 16th for R21 and R03, 25th for R15 awards to institutions with little NIH funding). We placed the within-investigator excess before each new-application due date in proportion to the number of funded applications on that schedule and fitted the sum to the national week-by-week profile with a single amplitude. The prediction has no free timing. It tracked the observed profile, explaining a share of 0.63 of the week-to-week variation on bioRxiv and 0.45 on medRxiv (weighted *R*^2^; Fig. 3A and table S13). Moving the surges away from the due dates lowered the bioRxiv fit to near zero at four weeks (Fig. 3B and table S14). A smooth wave with freely fitted timing (a sinusoid) follows the shape more closely (*R*^2^ 0.77 and 0.65; text S4), but the surges take their timing from the due dates alone and still put the bioRxiv peak within a week of where it is observed. Most of the remaining misfit is the seasonal dip in the fellowship window (Fig. 1D). On medRxiv the fit is best with the surges placed one to two weeks earlier, as expected if authors leave a margin for its longer screening (13, 14) (text S4).

**Fig. 3.**
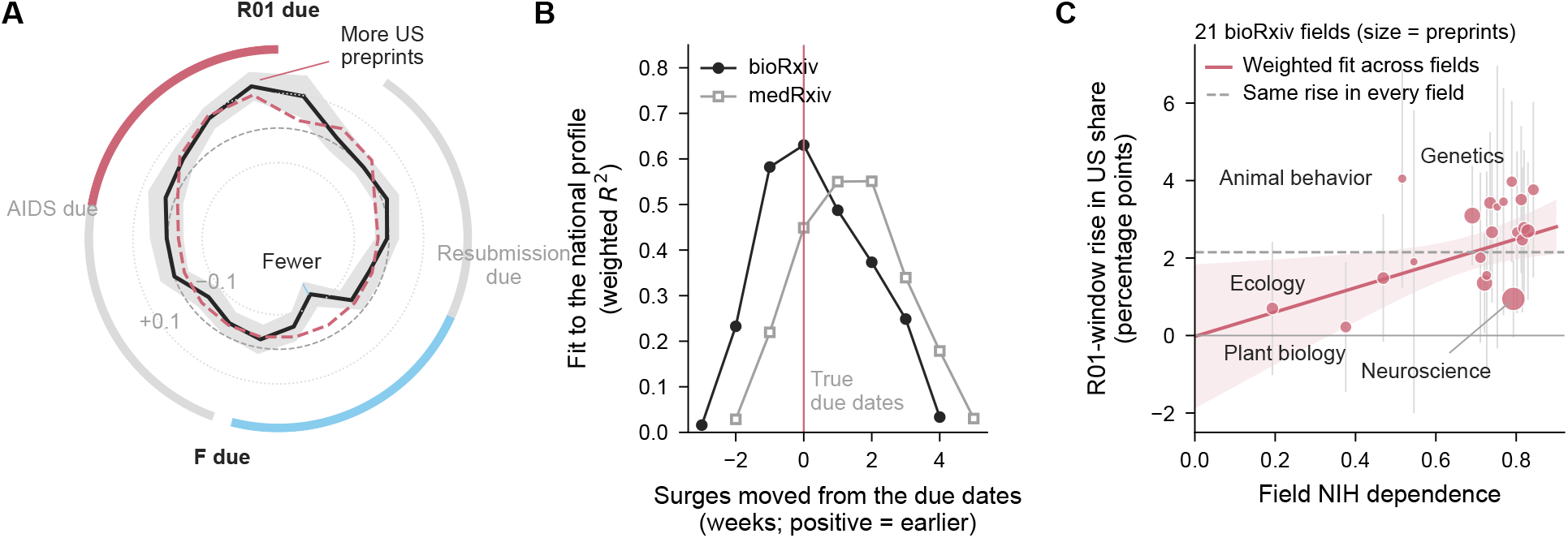
One surge, repeated on every due date, places the national peak. (**A**) The four-month cycle drawn as a clock, with the R01 due date at the top and time running clockwise. The radius is the log ratio of US to non-US daily first versions on bioRxiv (Poisson model with day fixed effects, which compares US with non-US preprints on the same day), relative to the dashed circle of no deviation; dotted circles, *±*0.1; bands, 95% confidence intervals. The dashed rose curve places the individual surge (the profile of Fig. 2B in weekly bins) before every new-application due date, weighted by the number of applications, with its height (amplitude) fitted and its timing fixed by the due dates. (**B**) How well that prediction fits the national bioRxiv (black) and medRxiv (gray) profiles (weighted *R*^2^) when the surges are moved away from the due dates. Fits that would need a negative height (amplitude), which would mean fewer US preprints before the due dates, are not shown. (**C**) R01-window rise in the US share of first versions for 21 bioRxiv fields against the share of funded US preprints in the field that acknowledge NIH (NIH dependence). Marker area is proportional to the number of preprints; the rose line and band are the inverse-variance weighted fit and its 95% confidence interval. The dashed line is what a calendar shared by all fields would predict, the same rise everywhere. The five categories with the least dependence are ecology, plant biology, evolutionary biology, animal behavior and cognition, and synthetic biology.

Because the amplitude is fitted, the match concerns when the national peak falls. How much of the rise applicants carry follows from arithmetic and one assumption. Funded applications are about one in five of those submitted (15), and our links reach 81% of funded investigators, so linked investigators hold about one application in six. They supply 5% to 25% of the national rise, so the national rise is four to twenty times theirs. All applicants, six times as many, would produce about that much if unfunded applicants, who face the same date and the same reason to post, responded as funded ones do. We assume they do. Failed first submissions, the unfunded applications we can see, show about 0.6 of the funded effect (0.052 against 0.087; text S3), so the calculation is an upper bound. The daily counts of authors outside the linked sample, who include unfunded applicants and investigators we could not link, rose at least as much as those of linked ones (0.173 against 0.134 on the log scale; text S4). Resubmitters carry the full surge individually (Fig. 2C). Nationally their date falls on the seasonal descent toward the fellowship trough. The drop there was 0.035 (SE 0.052; table S15 and text S4), an estimate too noisy to detect a drop half as large as the one at the R01 date.

Across 21 bioRxiv subject categories, the R01-window rise in US share grew with the share of funded US preprints that acknowledged NIH, by 3.14 percentage points per unit of dependence, a share from 0 to 1 (SE 1.19), with an intercept of *−*0.02 (SE 0.85) (Fig. 3C and table S16). The slope rests on the fields that depend least on NIH. The five categories with dependence below 0.6 rose by 1.19 points on average (SE 0.46) and the sixteen at 0.69 to 0.84 by 2.38 (SE 0.23), a difference of 1.19 (SE 0.51); on daily counts the difference was 0.031 on the log scale (SE 0.024). Ecology and plant biology, the two least dependent, barely moved (+0.70, SE 0.88, and +0.22, SE 0.85), but animal behavior and cognition, at 0.52, rose by 4.05 (SE 1.44), and neuroscience, the largest category and among the most dependent (0.79), rose by 0.95 (SE 0.51). The fields therefore show a smaller rise where dependence on NIH is lowest and no further rise with dependence among the more dependent categories (no dose response) (text S1). The gradient held with group-by-calendar-month fixed effects that absorb each field’s own season, such as summer field work (0.21 on daily counts, SE 0.09, against 0.10, SE 0.05, without them; table S8).

### What is released early is the science of the application

Preprints released by the surge should read like the application, and the resemblance should begin when the surge begins. If the deadline sets only when work appears, they should otherwise look like the investigator’s other preprints.

We compared the 763 last-author preprints posted in the 14 days before an investigator’s own due date with the same investigator’s other preprints, with fixed effects for investigator and month. Text similarity between a preprint and the abstract of the award that followed, measured as the overlap in distinctive vocabulary on a scale from 0 to 1, was 0.035 higher (SE 0.008) on a base of 0.162, whereas similarity to the investigator’s other awards did not change (0.000, SE 0.004) (Fig. 4B). The similarity rose only in the final two weeks. Relative to preprints posted six months to a year before the due date, it was 0.037 higher (SE 0.008) in the last 14 days and no more than 0.007 higher at any earlier interval (Fig. 4A). The rise in content and the rise in posting fall in the same two weeks, and adjusting for distance to the due date left a difference of 0.029 (SE 0.010). Restricted to preprints posted within 90 days of the due date the difference was 0.028 (SE 0.013), and within 45 days 0.021 (SE 0.020), so part of the resemblance reflects closeness in time to the application (table S17).

**Fig. 4.**
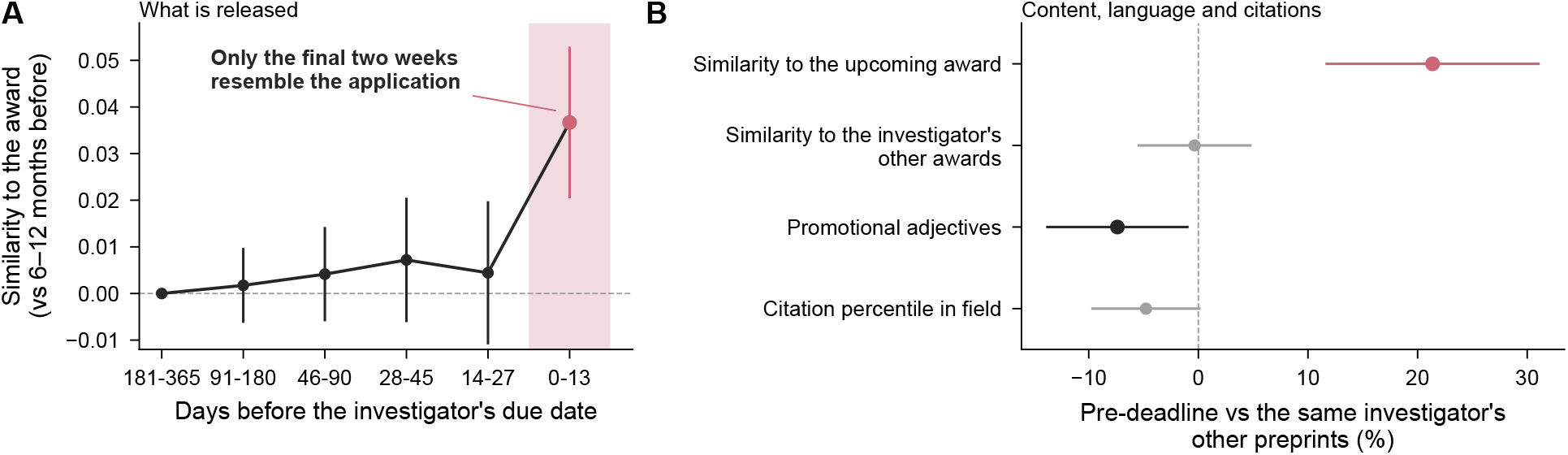
What is released early is the science of the application. (**A**) Text similarity (overlap in distinctive vocabulary) between an investigator’s preprint and the abstract of the award that followed, by days before the investigator’s due date, relative to preprints posted 181 to 365 days before, comparing each investigator’s preprints with one another and adjusting for posting month (investigator and month fixed effects). Only preprints posted in the final two weeks resemble the application more, the same two weeks as the posting excess in Fig. 2. Bins are of unequal width and drawn at equal spacing. (**B**) Preprints posted in the 14 days before the investigator’s own due date compared with the same investigator’s other preprints, as a percentage of the comparison mean. Similarity to the forthcoming award rises, and similarity to the investigator’s other awards does not. These preprints use no more of the promotional adjectives on a published list built from grant abstracts, and their citation rank among preprints of the same field and month (citation percentile) is slightly lower; fig. S3B shows all content measures. Rose and black mark estimates whose 95% confidence interval lies above or below zero, gray those that include it. Error bars, 95% confidence intervals.

These preprints were not written with more of the promotional adjectives that have spread through NIH grant abstracts. With the published list of 139 such adjectives (16), which has also been applied unchanged to journal abstracts (17), they had 0.83 fewer per 1,000 words than the same investigator’s other preprints (SE 0.36) on a base of 11.2, and 0.78 fewer (SE 0.35) on a base of 10.5 without “significant”, which in research abstracts mostly marks statistical significance (17). Without “significant” and adjusting for distance to the due date, the difference was *−*0.90 (SE 0.68), and its 95% confidence interval excludes an increase larger than about 4% of the base (tables S17 and S18). The list counts adjectives and does not measure claims that outrun the data.

Nor were these preprints better cited or more often published. Ranked by citations in the year of posting and the two that followed, among first versions of the same server, field and month, they stood 2.7 percentile points below the same investigator’s other preprints (SE 1.4; mean 55.4). Of those posted through 2023, 62% had reached a journal by June 2026, against 65% of the same investigators’ other preprints (within-investigator difference *−*3.4 points, SE 2.3; table S18 and text S4). NIH applicants thus release the science of their applications in the two weeks before NIH will judge them, and because thousands of labs work to the same due dates, their deadlines add up to a national rhythm in when science becomes public.

## Discussion

Funding deadlines act as a zeitgeber for science. Investigators’ output follows a slow cycle around their grants, higher around each application (table S19), as funding modestly raises output (18). NIH’s receipt calendar sets where in that cycle results become public. These data cannot tell whether the calendar adds preprints or only moves them, and the claim here concerns timing.

The consequence is that a funder’s calendar, as well as its awards, shapes the public record. Taking the non-US change as the baseline, about one in eleven US first versions on bioRxiv, and one in nine on medRxiv, posted in the four weeks before an R01 due date is there because of the deadline, several hundred preprints a year or 1.7% to 2.6% of US output from 2021 to 2025 (table S20). What these preprints carry is the science of the application, available to reviewers and competitors at the moment of submission. The deadline times it without selecting an investigator’s most cited work, and 38% of these preprints had not reached a journal at least two and a half years later, against 35% of the same investigators’ other preprints. Time series of US biomedical preprints carry this four-month rhythm, so analyses of trends, of seasonality or of policies dated near the due dates should account for it.

The evidence establishes the clock. The excess follows each investigator’s own date while the academic calendar stays fixed, the national ratio drops on the due date, and the individual surges place the national peak. How the clock couples to disclosure is less certain. Two mechanisms predict a peak that ends on the due date. Investigators may release results to make them citable, or the deadline may force the paper and the application to be finished from the same analyses. The evidence does not separate them. The citation rule is not shown to be necessary: no rhythm was detectable before 2017 in a comparison with little power, and the rhythm grew clearer only from 2021 (text S6 and table S1). The currency is supported in direction only: the excess appeared for R01 applications (0.095, SE 0.050) and not for R21 and R03 applications, which do not require preliminary data (*−*0.031, SE 0.075). The difference between the two cannot be distinguished from zero (text S3). We offer the currency and the exchange rule as predictions for future changes in funders’ rules.

The result returns to an old question. Merton described the institutions that govern when scientists disclose their work (1), and work on reactivity, people’s response to being measured, has shown how evaluation reshapes what people do (12). Here evaluation reshapes the timing of disclosure. Once an evaluation accepts a public product as evidence, the evaluation’s calendar can become a calendar of disclosure, as NIH’s has. Institutions that allocate resources can thereby govern when knowledge becomes available to everyone, although none set out to regulate disclosure.

Entrainment should occur wherever the three conditions meet. An evaluation with fixed dates, reviewers who value recent results and a rule that admits public products as evidence should pull disclosure toward its dates, and removing any one condition should weaken the pull. Deadlines are known to concentrate effort (19, 20), and the three conditions extend this to public disclosure across a national literature. They predict peaks of preprints before the submission deadlines of conferences that accept preprinted work and before research assessments that count outputs up to a fixed cutoff (census) date, and no peak where review is continuous or preprints cannot be cited. As funders move to require preprints (21, 22), more evaluations will meet these conditions. Social life is timed by institutional calendars (23), and the cycle of credibility that links papers, funding and new data (24) gives a funder’s calendar a place to act.

The study has limits. Country was assigned from the corresponding author, and non-US coauthors of US applicants are pulled toward the same dates, so both attenuate the contrast between US and non-US authors. Canadian authors, who can apply for R01 grants but not NIH fellowships, show part of the rhythm, so some of the national pattern may be shared with North American academic calendars (text S5). Posting dates trail submission by the servers’ screening, which is longer on medRxiv and blurs a drop at the due date there (13).

NIH can measure the pre-deadline peak at low cost. A cited preprint must carry its identifier, so NIH’s systems can read its posting date. A structured field for cited preprints would let NIH measure, for funded and unfunded applications alike, how much of the evidence it reviews was released in the final two weeks. Its rules exclude preprints from the post-submission update of preliminary data (25, 26), and admitting them for one grant type would show whether the peak follows the citability of results or the due date itself. Two changes NIH has already made are natural experiments it can evaluate with its own records. The removal of AIDS-designated due dates, effective for the January 2027 funding (council) round (27) and so for applications due from mid-2026, should move the posting peak of US HIV researchers about four weeks later, a test our aggregate data lack the power for (table S21). The end of continuous submission, which let frequent NIH reviewers apply at any time, with the last such applications accepted on 10 August 2026 (28), puts experienced investigators on the standard due dates, with a grace period of at most two weeks for recent reviewers, and their posting should acquire the pre-deadline peak.

## Materials and Methods

### Study design

The central prediction, that the US share of first versions rises before R01 due dates relative to the non-US share, with the fellowship and AIDS-designated schedules and the 2017 rule that let applications cite preprints as comparisons and shifted due dates as a null, was set before the data were downloaded. The within-investigator design, the discontinuity and weekend tests, the kernel model and the content analyses were developed during the analysis.

### Preprints

We downloaded metadata for all versions of bioRxiv preprints (January 2017 to October 2026) and medRxiv preprints (June 2019 to October 2026) from the servers’ application programming interface and used first versions: 345,602 on bioRxiv and 89,842 on medRxiv. Full-text JATS XML came from the servers’ text-and-data-mining repository on Amazon S3. The servers changed their DOI prefix from 10.1101 to 10.64898 on 1 December 2025 (8), and we joined records on the DOI suffix. We excluded preprints whose title or abstract mentioned COVID-19, SARS-CoV-2, coronavirus, 2019-nCoV, pandemic or lockdown. Bibliographic data, funders, award acknowledgments, journal versions and citations come from the OpenAlex snapshot of 26 June 2026 (29). The pre-2017 comparison also used bioRxiv metadata from November 2013 to December 2016.

### Country of the corresponding author

Each preprint was assigned to the country of its corresponding author using, in order, the country-code domain of the corresponding email (.edu and .gov as US), the Research Organization Registry match of the corresponding institution and the first affiliation country in the JATS XML. The main analyses use 315,824 bioRxiv and 60,718 medRxiv non-COVID first versions with a known country. Because email and JATS were unavailable before 2017, the pre-2017 comparison assigned country in both periods from the Research Organization Registry match of the corresponding institution alone.

### The NIH calendar

Each day was placed by its distance to the next R01 due date for new applications (5 February, 5 June, 5 October). We defined windows from 28 days before through the due date (29 days) for the R01, AIDS-designated (7 January, 7 May, 7 September), fellowship (8 April, 8 August, 8 December) and resubmission and renewal (5 March, 5 July, 5 November) due dates, which together cover 95% of the year. We used the published dates. A due date that falls on a weekend moves to the next business day (30), which moved 8 of the 30 R01 dates from February 2017 to October 2026 by one or two days. With business-day dates the discontinuity at the due date was nearly unchanged (table S22), and the weekend days in between behaved as days before the deadline (table S7). The 2020 extension (31) covered due dates from 9 March to 1 May and so no R01 date, and the 2025 extension (32) was announced on 24 November 2025, after applicants had planned for, and passed, the 5 October date, so neither moved a cue in advance. Near the R01 dates, Memorial Day falls one to two weeks before 5 June, which works against a drop, and Columbus Day can fall in the week after 5 October; no federal holiday is near 5 February, the cycle with the largest drop.

### Aggregate rhythm

For shares, we regressed an indicator of a US corresponding author on an indicator of one window at a time, with fixed effects for each four-month cycle and standard errors clustered by cycle. Per-group rate ratios came from Poisson models of each group’s daily counts with fixed effects for cycle and weekday. For comparisons between groups, we fitted Poisson models to daily counts of first versions by group (for example US and non-US), with fixed effects for day, group by cycle and group by weekday, and interactions of group with the window. Day fixed effects absorb everything common to all groups on a date. Week-by-week profiles replaced the window with indicators for 7-day bins of distance to the R01 due date, centered to average zero. Cycle-specific estimates restricted the panel to February, June or October cycles, and year-specific estimates to one calendar year. A seasonal check added group-by-calendar-month fixed effects to the daily-count model. Window length was varied to 21 (0 to 20 days) and 28 days (0 to 27). For the share model with the R01 and F windows together we report wild cluster bootstrap *P* values (Rademacher weights by cycle, 999 draws, restricted residuals). The drop at the due date was the US-by-indicator coefficient for the 7 days before the due date minus that for the 7 days from the due date on, in the daily-count model with day, group-by-cycle and group-by-weekday fixed effects and standard errors clustered by cycle.

Its null distribution repeated the estimate with every due date shifted by *s* days, 14 *≤ |s| ≤* 60 (94 shifts), and the rank *P* value is the share of shifts, counting the true dates, with a drop at least as large. Shifts of 28 to 31 days land on resubmission (later) and AIDS-designated (earlier) due dates. Without them in the null, the rank *P* value was 0.011 on both servers. Neighboring shifts overlap, so the shifts are fewer than 94 independent draws; measured against their spread, the bioRxiv drop lies 2.3 standard deviations above their mean. Group-by-day-of-year fixed effects would absorb the due date itself, which recurs on the same calendar day, and the business-day shifts are the variation such a model would leave. The holiday check used country codes to drop East Asian corresponding authors and the days from one day before to six days after each Lunar New Year and 1 to 7 October. The discontinuity analysis restricted the model to days within 7, 14 or 21 days of a due date (7 and 10 days at resubmission due dates, on the business day, and at a placebo date 45 days after the R01 date) and added US-specific linear trends in distance to the due date on each side; the drop is minus the US-by-after coefficient. Business-day due dates moved a date that fell on a weekend to the following Monday, and the weekend test entered indicators for the 7 days before the published date, the weekend days before the business day and the 7 days from the business day. The daily profile replaced the two indicators with one indicator per day from 21 days before to 20 days after the due date. The heatmap in Fig. 1A is the log ratio of US to non-US first versions in each year and 7-day bin, with 0.5 added to each count, centered within the year. For funder groups, US bioRxiv first versions were divided by their OpenAlex funders (NIH; NSF but not NIH; other funders only; none) and each group was compared with non-US first versions on the same day. The expected US count in each R01 window was the non-US count multiplied by the ratio of US to non-US counts outside the R01 and fellowship windows in the same year.

### Dependence on NIH

For each bioRxiv subject category, dependence was the share listing NIH among US non-COVID first versions with at least one OpenAlex funder. Per-category R01-window changes were regressed on dependence with inverse-variance weights. We report the weighted-least-squares standard error and, in table S16, the heteroskedasticity-robust (HC1) one. The two halves of the period are 2017–2021 and 2022–2026. We report leave-one-category-out ranges, count-based slopes from the Poisson model fitted per category, and the slope of the week-by-week profile on dependence for each 7-day bin, estimated the same way, and we report the intercept of that regression for each bin. The seasonal check fitted the per-category count model with group-by-calendar-month fixed effects, and the cycle check pooled per-cycle category estimates by inverse variance within each category before the regression. Institutional dependence was the leave-one-out NIH share among the funded preprints of the last author’s institution (US institutions with at least 30 funded preprints). Among US first versions, we regressed the dependence of the last author’s institution on the window indicator with field-by-cycle fixed effects, so the test asks whether preprints posted in the window come from more NIH-dependent institutions within the same field.

### The pre-2017 comparison

Periods were split at 24 March 2017, when NOT-OD-17-050 was announced (7); it covers applications due from 25 May 2017, and the first R01 due date after the split, 5 June 2017, is the first it covers. The later period runs through December 2019. We estimated window effects on the US share separately before and after, with cycle fixed effects and cycle-clustered standard errors, and their difference in a pooled model with a window-by-period interaction, for all categories and for categories with NIH dependence of at least 0.7 (dependence measured in 2017–2026).

### NIH awards and receipt cycles

From NIH RePORTER we retrieved 171,167 type 1 (new) and type 2 (renewal) awards with all principal investigators, funding opportunity announcements and spending categories. We used the first budget year of each competing segment and marked resubmissions by the A1 suffix. The due date of each application was inferred from the award notice month, which identifies the council round: notices in April or May were assigned the June due date of the previous year, notices in July or August the October due date of the previous year, and notices from October to December the February due date of the same year. The due day followed the activity code (R01 5th; R21, R03 and R34 16th; R15 25th; career awards 12th) (33); applications in the HIV/AIDS spending category used AIDS-designated dates. New applications used February, June and October dates, and resubmissions and renewals March, July and November dates, except R15 applications, which use the 25th of February, June and October for every application type. Awards made under requests for applications were excluded, as were notice months that map to more than one council round (January, February, March, June and September).

### Linking investigators to preprints

A principal investigator was linked to an OpenAlex author only when that author appeared, under the investigator’s family name and first initial, on a paper acknowledging one of the investigator’s own awards (institute code and serial number) among the 1.92 million OpenAlex works acknowledging NIH, and authors matching more than one investigator were dropped. Links by name and institution alone were not used (text S3). We took each linked author’s US last-author first versions. To rule out links made through the outcome, a sensitivity analysis anchored links only on works that are neither a bioRxiv or medRxiv preprint nor the journal version of one (141,438 of the 1.92 million works removed; table S9).

### Investigator-level estimates

For each investigator and each four-month cycle on the investigator’s schedule we counted preprints in the 28 days before the cycle’s due date (0 to 27 days) and in the rest of the cycle, restricted to cycles with at least one preprint. The within-investigator estimate regressed the window share of a cycle’s preprints on an indicator of the investigator’s own application cycle, with fixed effects for investigator and calendar cycle, weighted by the cycle’s preprint count and clustered by investigator, and we report it as a relative change, log(1 + *b/ȳ*), where *ȳ* is the weighted mean share, with standard error SE(*b*)*/ȳ*. Funded new applications were also split by activity code into R01 (with R37) and R21 or R03 applications, each with its own placebo distribution (100 draws). A further estimate replaced investigator fixed effects with investigator-by-cycle-family fixed effects (February, June and October schedules). The comparison across investigators used a Poisson model with investigator-by-cycle and calendar-cycle-by-window fixed effects, which estimates a log odds ratio of the window share. Estimates for the final two weeks replaced the 28-day window with a 14-day window. The profile compared, for investigators with preprints in both kinds of cycle, the share of a cycle’s preprints in each 14-day bin before the due date between application cycles and other cycles, with standard errors from 200 bootstrap resamples of investigators. Its placebo band replaced each investigator’s application cycle with one of the investigator’s other cycles, 200 times. The event study was a Poisson model of preprint counts in 14-day bins from 490 days before to 476 days after each due date, with event and calendar-bin fixed effects, separately for funded first submissions at the new-application due date and funded resubmissions at the resubmission due date. Two reference bins, 280 days before and after the due date, were needed because those fixed effects leave a linear trend in event time unidentified. Single-versus multi-investigator awards entered the comparison across investigators as an interaction with the treatment term.

A second within-investigator estimate dropped from the reference the cycles immediately before and after each application. Placebo distributions removed each investigator’s application cycles from the panel and assigned the same number of placebo cycles at random among the investigator’s remaining cycles, 200 times. Simulations held each investigator-cycle total fixed, drew the window count from a binomial distribution with the calendar-cycle share plus investigator-level noise, and added a known effect to application cycles (0 or 0.18 on the log-odds scale; 100 replicates). Three further designs used funded renewals; the renewal due date implied by the end of every R01 of a linked investigator, whether or not it was renewed; and the new-application due dates two and three cycles before each funded resubmission, both of which can hold the failed first submission. Total output per investigator-cycle was modeled with Poisson regression, investigator and calendar-cycle fixed effects and cycles with no preprints included (table S19). The share of US first versions written by linked investigators with an application due in that cycle, multiplied by the event-study effect, gave the part of the aggregate rise those investigators can supply.

### From the individual excess to the aggregate rhythm

The individual kernel was the within-investigator excess share in each of the 0 to 3 weeks before the investigator’s own due date, from the same profile in 7-day bins, with negative values, which do not differ from zero, set to zero so that the kernel describes an excess. Each new-application schedule (due on the 5th, 12th, 16th or 25th of February, June and October) received the kernel before its due date, weighted by the number of funded applications outside requests for applications on that schedule in RePORTER. The weighted sum, centered over the 18 weekly bins of the cycle, was fitted to the aggregate US-versus-non-US profile with one amplitude by inverse-variance weighted least squares. We report the correlation, the weighted *R*^2^ and the weighted residual sum of squares (*χ*^2^, 17 degrees of freedom) for new-application schedules alone, with AIDS-designated dates added, and with resubmission and renewal schedules added with equal weight (table S13); a further fit gave new-application and resubmission schedules separate non-negative amplitudes, with renewals dropped (table S15). For falsification the kernel was moved by *−*8 to +8 weeks and fitted at each shift, and fits with a negative amplitude, which invert the model, were not admissible. A sinusoid with a 126-day period and a free phase gave the descriptive benchmark (table S14). The coverage of the links was the share of investigators on funded new applications from fiscal year 2015 who were linked, and success rates came from the NIH Data Book (15).

### Content of pre-deadline preprints

Pre-deadline preprints were linked investigators’ last-author US first versions posted 0 to 13 days before the investigator’s own due date. They were compared with the same investigator’s other preprints in linear models with investigator and year-month fixed effects and standard errors clustered by investigator. Similarity was the cosine between term frequency–inverse document frequency vectors (sublinear term frequency, English stop words removed, terms in at least 3 and at most half of documents) of the preprint’s title and abstract and of the abstract of the award that followed, or the mean similarity to the investigator’s other award abstracts; the vocabulary was fitted on all compared preprints and award abstracts. Sensitivity analyses adjusted for the log of days to the due date or restricted the comparison to preprints posted within 90 or 45 days of it. The similarity profile regressed similarity on indicators of distance to the due date (0–13, 14–27, 28–45, 46–90 and 91–180 days; reference 181–365) with the same fixed effects. Promotional language was the rate per 1,000 abstract words of the 139 adjectives listed by Millar and colleagues (16), applied unchanged and, in a sensitivity analysis, without “significant” (17). Citation percentile counted citations to a first version and its journal version in the calendar year of posting and the two following years (OpenAlex counts by year) and ranked them within server, subject category and posting month, for first versions posted through 2023. Publication was a journal version linked by the server by June 2026. Similarity to the forthcoming award was split at its median for the comparison by similarity. Ratio estimates with 95% confidence intervals, *P* values and sample sizes were extracted from abstracts with regular expressions, and effect size was the absolute log of the first ratio. For shrinkage, journal abstracts were rebuilt from the OpenAlex inverted index, and shrinkage was the preprint’s absolute log ratio minus the journal version’s; it was compared between US and non-US preprints in the R01 window with week fixed effects. Citation crowding was tested by regressing log(1 + citations) on the R01 window interacted with field NIH dependence, with field-by-year and calendar-week fixed effects and standard errors clustered by week (table S23).

### Consistency checks

HIV preprints were those whose title, abstract or category matched HIV, AIDS, antiretroviral or PrEP. The HIV contrast was the interaction of US, HIV and the difference between AIDS and R01 windows in the Poisson day-count model, using AIDS-designated dates through 7 May 2026 (the last of these served a council round after which the dates were removed, which biases the contrast toward zero), and was repeated after removing the days from 21 before to 7 after each Conference on Retroviruses and Opportunistic Infections and IAS or International AIDS Conference. NIH intramural preprints had a corresponding author with an nih.gov address or an NIH institute affiliation. Minimum detectable effects are 2.8 standard errors (80% power, two-sided 5% level). Assumptions and full results are in text S1 and table S24.

### Data periods

Preprint metadata run to October 2026. Measures from OpenAlex (funders, award acknowledgments, journal versions and citations) run to the snapshot of 26 June 2026, and yearly estimates and counts use complete calendar years.

### Ethics

The study used public records of preprints, publications and NIH awards, involved no interaction with or intervention on any person and reports results only in aggregate, so it did not require review by an institutional review board.

### Software

Analyses used Python 3.12 with pandas, pyfixest and scikit-learn on Indiana University’s Quartz cluster, and DuckDB to query the OpenAlex snapshot.

## Supporting information

Data S1. Source data for all figure panels

## Acknowledgments

The authors acknowledge the Indiana University Pervasive Technology Institute for providing supercomputing and storage resources that have contributed to the research results reported within this paper (34). This research was supported in part by Lilly Endowment, Inc., through its support for the Indiana University Pervasive Technology Institute. The authors used Claude Fable 5.1 solely for copyediting and code verification, reviewed and revised the text as needed, and take full responsibility for its content. **Funding:** This study received no dedicated grant funding. **Author contributions:** H.W. conceived the study, collected the data, performed the analyses and wrote the paper. J.S. supervised the study and revised the paper. **Competing interests:** The authors declare that they have no competing interests. **Data and materials availability:** All data needed to evaluate the conclusions in the paper are present in the paper and/or the Supplementary Materials. All data are public: preprint records from the bioRxiv and medRxiv application programming interface, awards from NIH RePORTER, works and acknowledgments from OpenAlex and institution countries from the Research Organization Registry. Code that rebuilds every number, figure and table from these sources is available at https://github.com/Wang-Haining/zeitgeber.

## Supplementary Materials for

### Funding deadlines act as a zeitgeber for science

#### This PDF file includes

Supplementary Text (Texts S1 to S6) Figs. S1 to S3

Tables S1 to S35

#### Other Supplementary Materials for this manuscript include the following

Data S1

## Text S1. The drop at the due date and consistency checks: HIV researchers, NIH intramural scientists, fields, institutions and funders

**The drop at the due date.**

The drop is largest at the published date or one day after it on bioRxiv and two to three days after it on medRxiv, as expected from screening (text S4). It comes mostly from February: on bioRxiv it was 0.172 (SE 0.038) in February, 0.100 (SE 0.050) in June and 0.057 (SE 0.039) in October, and June and October pooled without February gave 0.074 (SE 0.031) (table S2). February is the cycle with no federal holiday near the due date (Materials and Methods). The fall is a discontinuity. Fitted with its own linear trend on each side of the due date, the bioRxiv ratio dropped by 0.106 (SE 0.048), 0.121 (SE 0.037) and 0.107 (SE 0.030) using the 7, 14 and 21 days on each side (bandwidths) (table S22). On medRxiv the break was not detectable. Lunar New Year and China’s National Day week lower posting in East Asia near the February and October due dates. Without East Asian authors in the comparison group and without those holiday weeks, the bioRxiv drop was 0.127 (SE 0.037) overall and 0.160 (SE 0.043) in February (table S25). A second rise 8 to 9 days after the R01 date (0.17 and 0.14), as high as the days just before the due date though below the peak, falls between the due dates of career awards (12th) and of R21 and R03 grants (16th) (Fig. 1B).

The main conclusions do not rest on the HIV or intramural case. Both are small (US HIV first versions about 0.5 per day on bioRxiv; NIH intramural about 0.8 per day), and the smallest effects their key contrasts could reliably detect (minimum detectable effects, 2.8 standard errors) are about 0.30 and 0.14 on the log scale. We state the assumptions under which each would identify the NIH effect.

**HIV researchers.**

Until 2026 NIH received AIDS-designated applications on 7 January, 7 May and 7 September, about four weeks before the standard R01 dates, so HIV researchers share the academic calendar of other US biomedical researchers but not their due date. The contrast asks whether US HIV researchers rise more in the AIDS window, relative to the R01 window, than the other groups do: the interaction of US, HIV and the difference between the AIDS and R01 windows in a Poisson model of daily counts for four groups (US HIV, US other, non-US HIV, non-US other) with day, group-by-cycle and group-by-weekday fixed effects (table S24). Three assumptions make it informative. (i) Parallel seasonality: without AIDS-designated dates, the seasonal profile of the US–non-US gap among HIV researchers would match that gap in other fields. (ii) Spillover can only attenuate: non-US HIV researchers who collaborate with US applicants can only be pulled toward the AIDS dates, which shrinks the contrast. (iii) Misclassification can only attenuate: preprints caught by the HIV keywords whose authors apply on standard dates dilute the contrast. Under (i) to (iii) the estimate is a lower bound. On bioRxiv the contrast was +0.09 (SE 0.11), and the AIDS-window term alone was +0.14 (SE 0.08), or +0.18 (SE 0.08) after removing days from 21 days before to 7 days after each Conference on Retroviruses and Opportunistic Infections and IAS or International AIDS Conference. US HIV researchers were also elevated in the R01 window, so the peak broadens as well as shifts. The end of AIDS-designated due dates gives a prospective test (main text).

**NIH intramural scientists.**

Intramural scientists do not apply on extramural due dates, but their papers pass through institutional clearance before posting, which adds a variable delay and would smooth a peak even if the incentive were present. We therefore use them descriptively. In the R01 window their daily counts rose by 0.077 on the log scale (SE 0.052), against 0.152 for US extramural and 0.056 for non-US authors on bioRxiv (difference from extramural *−*0.07, SE 0.05). On medRxiv the difference was +0.03 (SE 0.12).

**Fields.**

Dropping one bioRxiv category at a time left a slope of the R01-window share on NIH dependence between 2.38 (SE 1.10) and 4.23 (SE 0.80), and the slope was 3.87 (SE 0.77) in 2017–2021 and 1.81 (SE 1.60) in 2022–2026. On daily counts the slope was 0.10 on the log scale (SE 0.05; intercept 0.01). Among the sixteen categories with dependence of 0.69 to 0.84, neither the share change (Spearman correlation 0.30, *P* = 0.26) nor the count coefficient (0.32, *P* = 0.22) tracked dependence detectably. Estimated from the February and October cycles alone the count slope was 0.12 (SE 0.08), the same as from June alone (0.12, SE 0.13). Week by week, the gradient was 0.21 (SE 0.08) in the final week before the R01 due date and 0.21 (SE 0.09) in the week before the resubmission due date, and the intercept, the change expected in a field with no dependence on NIH, was *−*0.05 to 0.00 (SE 0.05 to 0.08) in the three weeks before the R01 due date (fig. S3A and table S26).

**Institutions and funders.**

Within a field, preprints from institutions more dependent on NIH did not rise more in the R01 window on bioRxiv (*z* = *−*0.07), although they did on medRxiv (*z* = 2.0). US authors acknowledging NIH, NSF but not NIH, other funders or no funder all rose in the R01 window relative to non-US authors, by amounts that could not be distinguished (table S27). Acknowledgments record past funding and not current applications, and most US biomedical laboratories apply to NIH whatever funded a given paper, so comparisons within the country cannot separate applicants from others. table S21 lists the remaining consistency checks with their predicted signs and minimum detectable effects, most larger than the rise itself.

**Fig. S1.**
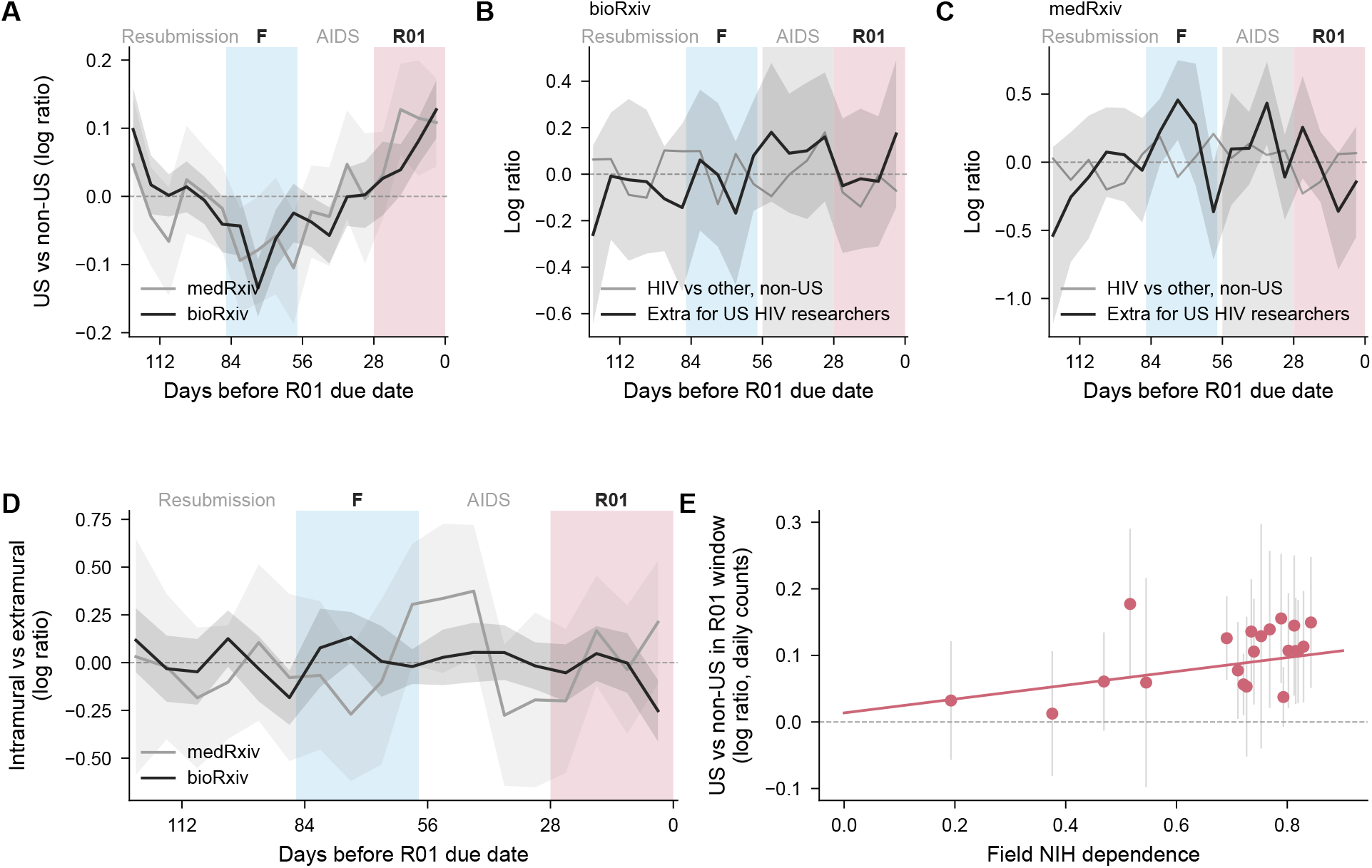
Week-by-week profiles on daily counts. (**A**) US versus non-US first versions (black, bioRxiv; gray, medRxiv). (**B**, **C**) HIV versus other fields among non-US authors (gray) and the additional US-specific term (black, with 95% confidence band), bioRxiv and medRxiv; the AIDS window is shaded gray. (**D**) NIH intramural versus US extramural first versions. (**E**) R01-window coefficient on daily counts by bioRxiv field against NIH dependence, with the inverse-variance weighted line. Poisson models with day, group-by-cycle and group-by-weekday fixed effects, 7-day bins shown relative to their average (centered to average zero). Rose and light blue shading mark the R01 and F windows. Bands and error bars, 95% confidence intervals.

## Text S2. Assumptions behind Eqs. 1 and 2

Equation 1 adds to each investigator’s own rate *h*_0_*_i_*(*t*) a multiplicative excess *c*(*t*) *A_i_ K*(*d_i_ − t*). The kernel *K* is positive in the two to three weeks before the due date and zero after it (Fig. 2B, C). The coupling *A_i_ ≥* 0 is larger for single-investigator awards (text S3). The exchange rule *c*(*t*) is 0 for due dates before preprints could be cited (25 May 2017) and 1 after. For Eq. 2, let US authors in field *f* hold a share *S_f_* of first versions, let a fraction *D_f_* of them depend on the funder, and let these apply on due dates with density *p_d_*. If the excess is small, it raises US output at cycle phase *ϕ* by a factor of about 1 + *Ā D_f_* (*K ∗ p_d_*)(*ϕ*) and leaves non-US output unchanged, so the US share changes by about *S_f_* (1 *− S_f_*) *Ā D_f_* (*K ∗ p_d_*)(*ϕ*). The constant *S_f_* (1 *− S_f_*) is absorbed into *Ā* in Eq. 2.

The model makes six statements that the data test. The rise grows with *D_f_* and is zero when *D_f_* = 0; the fields are consistent with the second statement and do not show the first within the dependent fields (Fig. 3C). At the individual level the excess is confined to the weeks in which *K* is positive (its support) (Fig. 2B, C; Fig. 4A). Because *K* is zero after the due date, the rate drops on the due date itself, which a smooth seasonal term in *h*_0_*_i_* does not produce (Fig. 1B, C). No rhythm was detectable while *c* = 0, in a comparison with little power (fig. S2). The rise in the aggregate is broader than the individual kernel because it convolves *K* with the spread of due dates across grant types (activity codes), summing the bump over all of them (Fig. 3A, B and tables S13 and S14). Because *h*_0_*_i_* varies with the grant cycle (table S19), the model separates the timing of release, which the deadline sets, from the volume of work, which it may not.

## Text S3. Investigator-level estimand, placebo and simulation

**Two comparisons (estimands).**

With fixed effects for investigator by cycle and for calendar cycle by window, a Poisson model of window and non-window counts compares an application cycle’s pre-deadline split with the calendar-cycle average across all investigators. This comparison across investigators gives a log odds ratio of 0.178 (SE 0.042) for funded new applications. On the scale of shares it is 0.130. Restricted to investigators observed in at least two cycles it is 0.114, and with investigator fixed effects, which make each investigator’s other cycles the reference, 0.087 (SE 0.039) (table S28).

**Placebo.**

A placebo cycle is a non-application cycle treated as if it were the application cycle. Placebo cycles are drawn only among each investigator’s non-application cycles. A draw among all of an investigator’s cycles would often return the true application cycle, because an investigator with *k* informative cycles and one application re-selects it with probability 1*/k*, and the placebo distribution would then carry part of the real effect. Drawn among non-application cycles, the placebo centers on zero in every design for both estimands (table S29).

**Simulation.**

On the observed panel of funded new applications we held each investigator-cycle total fixed, drew the window count from a binomial distribution whose log odds were the calendar-cycle share plus investigator noise (SD 0.5), and added a known effect of 0 or 0.18 to application cycles. Both estimators were unbiased when no effect was present (mean *−*0.002 and 0.000), with standard errors that matched the spread of the estimates and 95% intervals that contained the true value in 93% to 96% of replicates (coverage of 0.93 to 0.96) (table S30). With an effect of 0.18 the across-investigator estimator returned 0.171 and the within-investigator estimator 0.125, close to the 0.132 that a log-odds effect of 0.18 implies on the share scale.

**Further investigator-level estimates.**

Restricted to the final 14 days, the within-investigator increase was 12% (SE 6%), and the third week before the due date was 0.9 points higher in application cycles. Placebo estimates had mean 0.000 and SD 0.040, and for failed first submissions the placebo *P* was 0.045. By annual cycle the effect was +0.24 in February (SE 0.12), +0.02 in June (SE 0.08) and +0.08 in October (SE 0.06), largest where the aggregate drop at the due date is largest. The share of preprints posted before the next cycle’s due date was lower than in other cycles (*−*0.10, SE 0.04; table S29), as expected if reference cycles contain unobserved applications and the cycle right after a submission is least likely to contain another. Dropping the cycles adjacent to each application from the reference left the estimate at 0.079 (SE 0.040; table S31). The renewal clock of every R01, the due date implied by its scheduled end whether or not it was renewed, gave +0.03 (SE 0.03), and the difference between new applications and renewals was 0.04 (SE 0.06). Across investigators the effect was stronger for single-investigator than for multi-investigator awards (interaction *−*0.16, SE 0.08).

**Event study and design checks.**

Pooling first submissions and resubmissions, only the 14 days before the due date departed from the reference (+0.28 on the log scale, SE 0.06), and counts did not dip after submission. Failed first submissions, which were not funded, gave 0.052 (SE 0.033), but across investigators their estimate for the preceding cycle was as large (0.082 against 0.080), so this design does not show timing specific to the cycle (table S29). By activity code the estimates point the way the currency predicts. The excess was present for R01 applications, which reviewers judge on preliminary data (0.095, SE 0.050), and not detectable for R21 and R03 applications, which do not require them (*−*0.031, SE 0.075), or for funded renewals, which reviewers judge on progress (+0.04, SE 0.05). Neither difference can be distinguished from zero (tables S29 and S32).

**Resubmissions.**

Restricted to funded resubmissions (A1), the comparison across investigators within the same calendar cycle gave 0.197 (SE 0.066; 9,052 investigators, 382 pre-deadline preprints), against 0.168 (SE 0.053) for first submissions (A0). Text S4 describes the national profile at the resubmission dates.

**Inferred due dates.**

The due date of each application was inferred from the award notice month. If the mapping were wrong by one cycle for a given notice month, the effect would appear in the cycle before or after. By notice month, the estimate was largest in the inferred own cycle in four of six months. In May and August the cycle before was as large or larger, with overlapping intervals (table S12).

**Linkage.**

Investigators were linked to OpenAlex authors only through papers acknowledging their own awards. A preprint, or its journal version, that acknowledged the award could make its own author linkable and so select on the outcome. With anchors restricted to works that are neither, the within-investigator estimate was 0.087 (SE 0.039) and the failed first submissions gave 0.056 (SE 0.033) (table S9). Links by name and institution alone were confirmed by this anchor for 88% of investigators overall and 74% in a sample of 50 from each of four strata of surname frequency and identifier count, from 98% for uncommon surnames with a single OpenAlex identifier to 54% for common surnames with several (table S33). These links were not used.

## Text S4. Output, scale and consequences

Investigators’ total output was higher in the cycle before, the cycle of and the cycle after an application than in other cycles for most designs, by up to 26% (table S19). For the renewal clock, set by the project end date years in advance, output was equally elevated in the due cycle and the cycle after. We read this as the productivity cycle of a grant. The aggregate excess of US first versions in R01 windows grew from fewer than 200 a year before 2021 to 324 to 559 a year from 2021 to 2025, and the F windows held 137 to 385 fewer than expected each year from 2018 to 2025 (table S20), a deficit that calendar-month fixed effects attribute to season (table S8). At the field level, R01-window preprints in fields more dependent on NIH were cited no less than preprints posted at other times, including among non-US authors posting into the same weeks. General crowding in a field’s week did carry a citation cost (table S23). Within investigators and adjusting for distance to the due date, journal versions of pre-deadline preprints acknowledged the forthcoming award 6.0 percentage points more often (SE 4.4).

**How much applicants can supply.**

Linked investigators with an application due in a given cycle wrote 1.25% of US preprints, enough to supply 5% to 25% of the aggregate rise, and the links cover 81% of investigators on funded new applications (53,361 of 65,816). NIH funded 22% of R01-equivalent applications in fiscal year 2023 and 19% in 2024, so each funded application stands for about five submitted ones. If unfunded applicants posted as funded ones do, applicants could supply most of the rise. Failed first submissions show about 0.6 of the funded effect (0.052 against 0.087), and one applicant may submit several applications, so the estimate is an upper bound. US preprints whose last author was not a linked investigator rose at least as much as those whose last author was (0.173 and 0.134 on the log scale), consistent with trainees, unfunded applicants and unlinked investigators who face the same dates.

**Shape of the national profile.**

The kernel prediction correlated with the observed profile at 0.78 on bioRxiv and 0.66 on medRxiv, and its residuals exceed sampling error (*χ*^2^ = 52 on 17 degrees of freedom on bioRxiv; table S13). Moved one week earlier the bioRxiv fit was 0.49 and one week later 0.58, and on medRxiv the best fit, with the kernel one to two weeks earlier, was 0.55. A smooth wave with the period of the cycle and freely fitted timing (a sinusoid with a free phase) fitted better (*R*^2^ 0.77 and 0.65), peaking about 6 and 10 days before the R01 due date (table S14). The earlier medRxiv peak matches the later medRxiv drop at the due date (Fig. 1C): by the servers’ own accounts, medRxiv screening usually takes two to four days and longer when its clinical screeners correspond with the authors, against usually under 72 hours on bioRxiv (sources in the main text), so authors who need a preprint public by the due date would submit earlier, and those who submit close to it would appear after it.

**Resubmission dates.**

Resubmissions are due 28 to 31 days after new applications, so their 28-day window is the month after the R01 date. With a linear trend on each side, the national drop at the resubmission date was 0.035 (SE 0.052) with a 7-day bandwidth and 0.080 (SE 0.043) with 10 days, against 0.018 (SE 0.052) and 0.060 (SE 0.045) at a placebo date 45 days after the R01 date and 0.100 (SE 0.048) and 0.150 (SE 0.049) at the R01 date. By month, only November showed a detectable drop, with the 10-day bandwidth (0.120, SE 0.052) and not with 7 days (0.037, SE 0.080), and weekend days before a resubmission date that fell on a weekend did not behave like days before a deadline (*−*0.063, SE 0.053, on bioRxiv) (table S15). Without trends the drop statistic was 0.096 (SE 0.027) at 29 days after the R01 date, but it reached 0.085 (SE 0.028) at 45 days, where no NIH due date falls, as the profile descends toward the fellowship-window trough. Placed before the resubmission dates with a height (amplitude) of their own, the individual surges carry 0.15 of the amplitude per funded new application on bioRxiv and 0.02 on medRxiv, and they barely change the fit, whereas resubmission and renewal schedules added with the same weight as new applications lowered it to *R*^2^ = 0.22 and 0.10.

**Content.**

The number of versions of pre-deadline preprints (difference *−*0.003, SE 0.027), journal publication within 24 months (*−*3.2 percentage points, SE 2.3) and time to publication (+0.057 on the log scale, SE 0.051) did not differ from the same investigator’s other preprints. In citation percentile, the 346 of 556 that resembled the forthcoming award more than the median compared preprint ranked like the investigator’s other work (+0.4, SE 2.8), and the other 210 ranked lower (*−*8.0, SE 2.7). Those published were cited like the investigator’s other published work (*−*0.5, SE 1.5), and those not published were cited less (*−*4.3, SE 2.3) (table S18). Only 63 reported a ratio estimate, and their effect sizes changed sign with the specification (table S17). Across all preprints, the shrinkage of effect sizes between preprint and journal version did not differ detectably between the R01 window and other times (medRxiv, *−*0.020 on the log scale, SE 0.028, which allows up to about 0.04 more shrinkage in the window).

## Text S5. Other countries and funders

Canadian authors, who may apply for R01 grants but not for NIH fellowships, showed part of the rhythm: their share fell in the F window on bioRxiv and rose in the R01 window on medRxiv (table S34). Some of the aggregate pattern may therefore be shared across North American institutions. Before the annual deadlines of the National Natural Science Foundation of China (20 March, verified for 2019, 2020, 2022, 2023, 2025 and 2026) and of KAKENHI in Japan (verified due dates of 2022 to 2025), the share of authors from those countries among non-US first versions rose by 6.7% and 8.9% (relative). The same contrast at placebo dates 91 and 182 days before and after each due date was smaller in every case, and *z* for the true dates was below 2. Neither funder asks for recent preliminary data in the way NIH does, and we regard these results as inconclusive.

## Text S6. Preprint citation in 2017

NIH announced on 24 March 2017 that applications due from 25 May 2017 could cite preprints. With bioRxiv first versions from November 2013 and country assigned in both periods from the ROR match of the corresponding institution, no rhythm was detectable before the notice (compared with the period through December 2019), in all fields or in the fields most dependent on NIH (fig. S2, table S35). The contrast is underpowered: before 2017 there are 8,290 first versions in 11 cycles, the standard error of the R01-window effect is 1.38 points, and the minimum detectable effect is about 3.9 points.

**Fig. S2.**
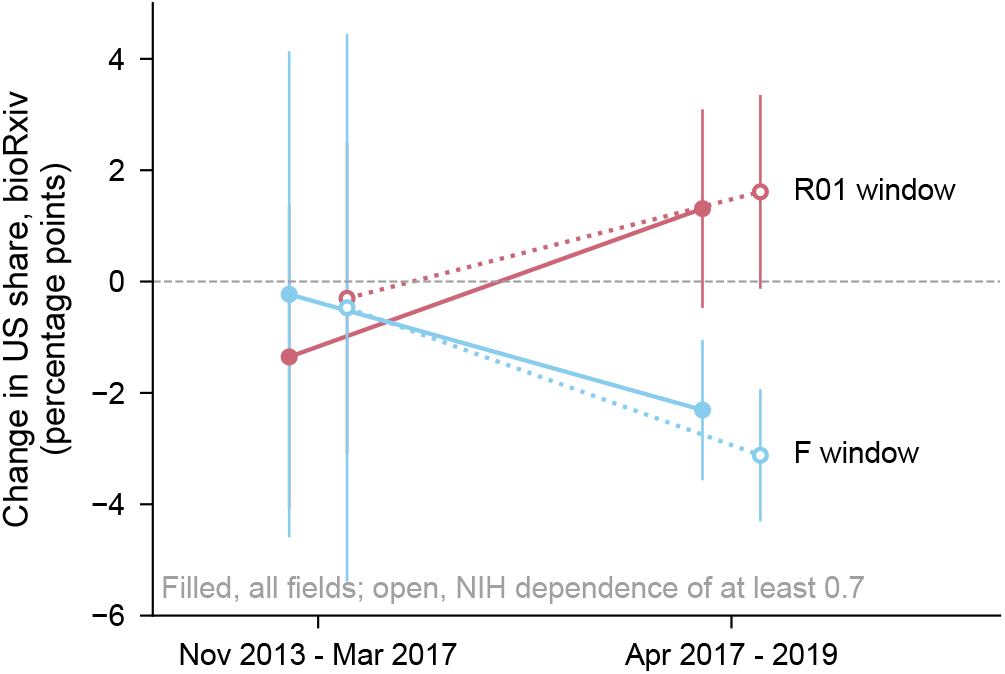
Before and after NIH allowed applications to cite preprints. (**2017**). Change in the US share of bioRxiv first versions in the R01 (rose) and F (light blue) windows, for all fields (filled) and fields with NIH dependence of at least 0.7 (open). Error bars, 95% confidence intervals.

**Fig. S3.**
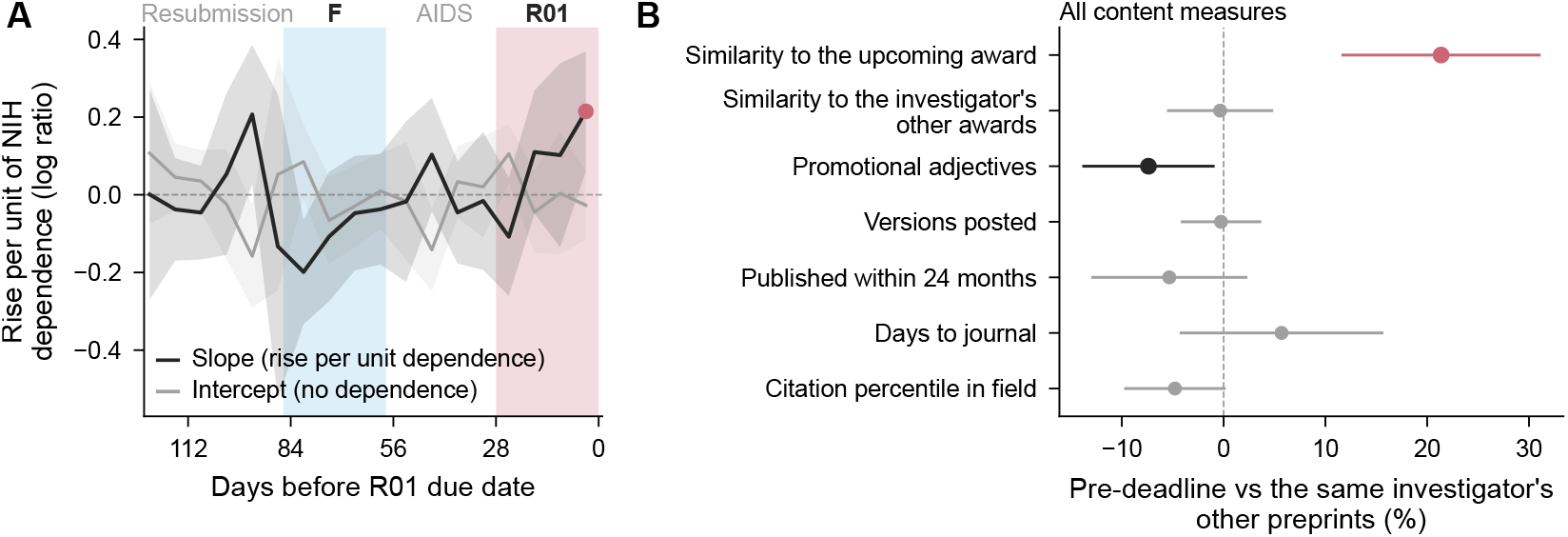
Fields week by week and all content measures. (**A**) For each week, how much more fields rise per unit of NIH dependence (slope, black) and the rise expected in a field with no dependence (intercept, gray), from field coefficients on daily counts. The slope is clearly positive in the final week before the R01 due date and in the week before the resubmission due date, and the intercept is near zero in the three weeks before the R01 due date. (**B**) All content measures of preprints posted in the 14 days before the investigator’s own due date, compared with the same investigator’s other preprints, as a percentage of the comparison mean (for logged outcomes, 100 times the log difference). Rose and black mark estimates whose 95% confidence interval lies above or below zero, gray those that include it. Error bars and bands, 95% confidence intervals.

**Table S1.** Year by year: US *×* window on daily counts (Poisson, day fixed effects, SE by cycle).

| Server | Year | R01 window | F window | R01 – F |
| --- | --- | --- | --- | --- |
| bioRxiv | 2017 | 0.048 (0.065) | –0.069 (0.025) | 0.117 (0.070) |
| bioRxiv | 2018 | –0.051 (0.033) | –0.077 (0.063) | 0.026 (0.042) |
| bioRxiv | 2019 | 0.101 (0.071) | –0.129 (0.014) | 0.230 (0.086) |
| bioRxiv | 2020 | 0.006 (0.062) | –0.107 (0.011) | 0.113 (0.055) |
| bioRxiv | 2021 | 0.113 (0.022) | –0.029 (0.010) | 0.142 (0.021) |
| bioRxiv | 2022 | 0.103 (0.067) | –0.014 (0.038) | 0.117 (0.088) |
| bioRxiv | 2023 | 0.105 (0.033) | –0.030 (0.019) | 0.135 (0.029) |
| bioRxiv | 2024 | 0.110 (0.018) | –0.052 (0.025) | 0.162 (0.041) |
| bioRxiv | 2025 | 0.084 (0.025) | –0.076 (0.031) | 0.159 (0.054) |
| medRxiv | 2020 | 0.013 (0.015) | –0.129 (0.034) | 0.142 (0.024) |
| medRxiv | 2021 | 0.012 (0.027) | –0.144 (0.104) | 0.156 (0.085) |
| medRxiv | 2022 | 0.083 (0.050) | –0.179 (0.037) | 0.261 (0.078) |
| medRxiv | 2023 | 0.118 (0.079) | –0.154 (0.098) | 0.272 (0.077) |
| medRxiv | 2024 | 0.174 (0.065) | –0.025 (0.019) | 0.198 (0.063) |
| medRxiv | 2025 | –0.015 (0.049) | –0.006 (0.026) | –0.009 (0.044) |

**Table S2.** The drop at the R01 due date. Log ratio of US to non-US daily first versions in the 7 days before the due date minus the 7 days from the due date on (Poisson, day, group-by-cycle and group-by-weekday fixed effects, SE clustered by cycle). The null distribution is the same statistic with every due date shifted by 14 to 60 days in either direction (94 shifts); rank *P* is the share of shifts with a drop at least as large, counting the true date.

| Server | Cycles | Drop (SE) | Null mean | Null SD | Null max | Rank $P$ |
| --- | --- | --- | --- | --- | --- | --- |
| bioRxiv | February, June and October | 0.107 (0.031) | –0.004 | 0.047 | 0.096 | 0.011 |
| bioRxiv | February | 0.172 (0.038) | –0.009 | 0.081 | 0.125 | 0.011 |
| bioRxiv | June | 0.100 (0.050) | –0.011 | 0.066 | 0.190 | 0.063 |
| bioRxiv | October | 0.057 (0.039) | 0.009 | 0.075 | 0.200 | 0.242 |
| bioRxiv | June and October<br>(inverse-variance pooled) | 0.074 (0.031) | – | – | – | – |
| medRxiv | February, June and October | 0.096 (0.053) | –0.013 | 0.061 | 0.132 | 0.032 |
| medRxiv | February | 0.125 (0.069) | –0.023 | 0.110 | 0.243 | 0.095 |
| medRxiv | June | 0.085 (0.135) | –0.003 | 0.072 | 0.178 | 0.147 |
| medRxiv | October | 0.095 (0.073) | –0.014 | 0.113 | 0.181 | 0.168 |
| medRxiv | June and October<br>(inverse-variance pooled) | 0.093 (0.064) | – | – | – | – |

**Table S3.** Change in the US share of first versions in each window, estimated one window at a time against the rest of its four-month cycle (linear probability model, cycle fixed effects, SE clustered by cycle). These are the estimates cited in the main text; table S5 gives the R01 and F windows estimated jointly and on daily counts.

| Server | Window | US share, pp (SE) | Cycles |
| --- | --- | --- | --- |
| bioRxiv | R01 | 2.26 (0.33) | 30 |
| bioRxiv | AIDS | −0.80 (0.31) | 30 |
| bioRxiv | F | −1.89 (0.29) | 30 |
| bioRxiv | Resubmission | 0.46 (0.32) | 30 |
| medRxiv | R01 | 2.70 (0.53) | 22 |
| medRxiv | AIDS | 0.04 (0.52) | 22 |
| medRxiv | F | −2.46 (0.51) | 22 |
| medRxiv | Resubmission | −0.42 (0.55) | 22 |

**Table S4.** R01-window change in the US share of first versions (pp, cycle fixed effects, SE clustered by cycle) for windows of different length, counted in days before the due date.

| Server | Window | US share, pp (SE) |
| --- | --- | --- |
| bioRxiv | 21 days (0–20) | 2.55 (0.39) |
| bioRxiv | 28 days (0–27) | 2.32 (0.33) |
| bioRxiv | 29 days (0–28, main text) | 2.26 (0.33) |
| medRxiv | 21 days (0–20) | 3.30 (0.58) |
| medRxiv | 28 days (0–27) | 2.92 (0.51) |
| medRxiv | 29 days (0–28, main text) | 2.70 (0.53) |

**Table S5.** Aggregate rhythm on daily counts and the R01. *−* **F contrast.** Rate-ratio rows are log rate ratios from Poisson models of each group’s daily counts (cycle and weekday fixed effects). US-versus-non-US rows come from one Poisson model with both window indicators and day fixed effects. Share rows come from one linear model with both window indicators, which is why the R01 share differs from the one-window estimate in table S3. Wild cluster bootstrap: Rademacher weights by cycle, 999 draws, *P* for the null of zero.

| Server | Measure | Window | Estimate (SE) | Wild bootstrap $P$ |
| --- | --- | --- | --- | --- |
| bioRxiv | rate ratio, US | R01 | 0.150 (0.018) | – |
| bioRxiv | rate ratio, US | F | –0.065 (0.011) | – |
| bioRxiv | rate ratio, non-US | R01 | 0.056 (0.016) | – |
| bioRxiv | rate ratio, non-US | F | 0.013 (0.015) | – |
| medRxiv | rate ratio, US | R01 | 0.170 (0.024) | – |
| medRxiv | rate ratio, US | F | –0.085 (0.029) | – |
| medRxiv | rate ratio, non-US | R01 | 0.053 (0.019) | – |
| medRxiv | rate ratio, non-US | F | 0.024 (0.016) | – |
| bioRxiv | US vs non-US, day FE | R01 | 0.075 (0.014) | – |
| bioRxiv | US vs non-US, day FE | F | –0.053 (0.012) | – |
| bioRxiv | US vs non-US, day FE | R01 – F | 0.128 (0.017) | – |
| medRxiv | US vs non-US, day FE | R01 | 0.092 (0.023) | – |
| medRxiv | US vs non-US, day FE | F | –0.078 (0.023) | – |
| medRxiv | US vs non-US, day FE | R01 – F | 0.170 (0.031) | – |
| bioRxiv | US share (pp), 30 clusters | R01 | 1.86 (0.33) | 0.000 |
| bioRxiv | US share (pp), 30 clusters | F | –1.28 (0.29) | 0.000 |
| bioRxiv | US share (pp), 30 clusters | R01 – F | 3.13 (0.39) | 0.000 |
| medRxiv | US share (pp), 22 clusters | R01 | 2.14 (0.52) | 0.002 |
| medRxiv | US share (pp), 22 clusters | F | –1.75 (0.49) | 0.002 |
| medRxiv | US share (pp), 22 clusters | R01 – F | 3.89 (0.69) | 0.000 |

**Table S6.** Rhythm by annual cycle. US *×* window coefficients on log daily counts (Poisson, day fixed effects, SE clustered by cycle), and the bioRxiv field gradient of the R01-window coefficient on NIH dependence (21 categories, inverse-variance weighted, HC1 SE).

| Server | Cycle | US × R01 | US × F | Gradient slope | Gradient intercept |
| --- | --- | --- | --- | --- | --- |
| bioRxiv | February | 0.040 (0.015) | –0.076 (0.015) | 0.152 (0.105) | –0.079 (0.052) |
| bioRxiv | June | 0.165 (0.016) | –0.119 (0.016) | 0.124 (0.129) | 0.087 (0.087) |
| bioRxiv | October | 0.063 (0.014) | –0.038 (0.019) | 0.032 (0.047) | 0.028 (0.035) |
| medRxiv | February | 0.085 (0.043) | –0.085 (0.022) | – | – |
| medRxiv | June | 0.096 (0.050) | –0.128 (0.053) | – | – |
| medRxiv | October | 0.160 (0.035) | –0.113 (0.044) | – | – |

**Table S7.**
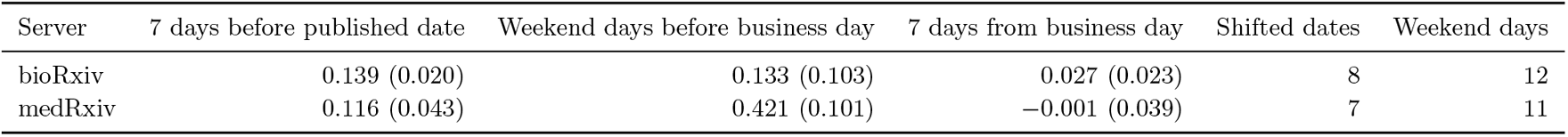
When the due date falls on a weekend. US *×* indicator coefficients on daily counts for the 7 days before the published date, the weekend days between the published date and the business day on which NIH accepts applications, and the 7 days from that business day on (same model as table S22). If posting follows the date NIH uses, the weekend days resemble the days before the deadline.

**Table S8.** Annual season. Upper rows: US *×* window on bioRxiv daily counts, and the slope of the per-field window coefficient on NIH dependence (21 fields, inverse-variance weights, HC1 SE), without and with group-by-calendar-month fixed effects. Lower rows: the R01-window slope from the per-cycle field estimates, pooled by inverse variance within field over the cycles named.

| Window | Month FE | All fields (SE) | Slope on dependence (SE) | Intercept (SE) |
| --- | --- | --- | --- | --- |
| R01 | no | 0.092 (0.014) | 0.104 (0.050) | 0.014 (–) |
| R01 | yes | 0.073 (0.021) | 0.210 (0.086) | –0.077 (–) |
| F | no | –0.078 (0.012) | –0.122 (0.047) | 0.006 (–) |
| F | yes | –0.014 (0.015) | –0.085 (0.043) | 0.051 (–) |
| R01, February and October cycles | no | – | 0.123 (0.083) | –0.049 (0.052) |
| R01, all cycles | no | – | 0.136 (0.095) | –0.023 (0.057) |
| R01, June cycle | no | – | 0.124 (0.129) | 0.087 (0.087) |

**Table S9.** Investigator-level estimates with links anchored only on works that are neither a bioRxiv or medRxiv preprint nor the journal version of one (141,438 of 1,918,958 acknowledging works removed; 66,364 investigators linked). Placebo: 100 draws among non-application cycles.

| Design | Estimand | Estimate (SE) | Placebo 97.5% | Placebo $P$ | Investigators | Pre-deadline preprints |
| --- | --- | --- | --- | --- | --- | --- |
| Funded new applications | Across investigators | 0.175 (0.042) | 0.057 | 0.010 | 15,552 | 981 |
| Funded new applications | Within investigator | 0.087 (0.039) | 0.049 | 0.010 | 15,552 | 981 |
| Failed first submission | Across investigators | 0.081 (0.037) | 0.052 | 0.010 | 13,193 | 1,238 |
| Failed first submission | Within investigator | 0.056 (0.033) | 0.081 | 0.089 | 13,193 | 1,238 |

**Table S10.** Within-investigator estimate for funded new applications with investigator fixed effects and with investigator-by-cycle-family fixed effects (February, June and October schedules), which compare an application cycle only with the investigator’s other cycles of the same annual family. Placebo: 100 draws among non-application cycles.

| Fixed effects | Log ratio (SE) | Share, pp (SE) | Placebo $P$ | Investigators |
| --- | --- | --- | --- | --- |
| Investigator | 0.087 (0.039) | 2.28 (0.98) | – | 15,906 |
| Investigator × cycle family | 0.090 (0.047) | 2.36 (1.20) | 0.069 | 15,906 |

**Table S11.**
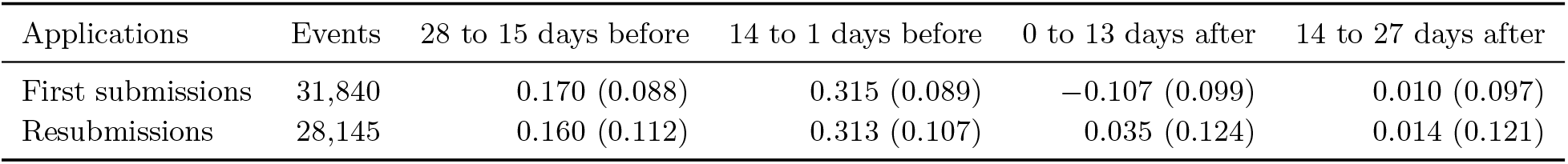
Linked investigators’ preprint counts around their own due date: funded first submissions at the new-application date and funded resubmissions at the resubmission date one month later. Poisson model of counts in 14-day bins with event and calendar-bin fixed effects, SE clustered by investigator (as Fig. 2C).

| Applications | Events | 28 to 15 days before | 14 to 1 days before | 0 to 13 days after | 14 to 27 days after |
| --- | --- | --- | --- | --- | --- |
| First submissions | 31,840 | 0.170 (0.088) | 0.315 (0.089) | –0.107 (0.099) | 0.010 (0.097) |
| Resubmissions | 28,145 | 0.160 (0.112) | 0.313 (0.107) | 0.035 (0.124) | 0.014 (0.121) |

**Table S12.** Within-investigator estimates by award notice month (checks the inferred due date: a mapping wrong by one cycle would move the effect to the cycle before or after) and by annual cycle.

| Subset | Cycle before | Own cycle | Cycle after | Pre-deadline posts, own cycle |
| --- | --- | --- | --- | --- |
| Notice month 4 | 0.037 (0.110) | 0.126 (0.101) | -0.058 (0.098) | 139 |
| Notice month 5 | 0.099 (0.108) | -0.013 (0.091) | -0.149 (0.085) | 157 |
| Notice month 7 | -0.110 (0.074) | 0.031 (0.079) | -0.023 (0.079) | 241 |
| Notice month 8 | 0.109 (0.078) | 0.085 (0.076) | -0.109 (0.069) | 300 |
| Notice month 10 | 0.854 (1.778) | 0.866 (0.907) | 0.292 (1.196) | 5 |
| Notice month 11 | 0.101 (0.195) | 0.128 (0.219) | -0.104 (0.186) | 49 |
| Notice month 12 | -0.171 (0.121) | 0.168 (0.126) | -0.025 (0.119) | 114 |
| Cycle family February | - | 0.242 (0.121) | - | 168 |
| Cycle family June | - | 0.022 (0.084) | - | 295 |
| Cycle family October | - | 0.083 (0.064) | - | 540 |

**Table S13.** The individual surge (investigator kernel) fitted to the national (aggregate) rhythm. The excess in the 0–3 weeks before an investigator’s own due date (Fig. 2B) is placed before the due date of each schedule, weighted by the number of funded applications on that schedule, and fitted to the aggregate week-by-week profile with one amplitude (in units of 10*^−^*^4^ on the log ratio per weighted application).

| Server | Schedules | Correlation | Weighted $R^2$ | Amplitude | $\chi^2$ | df |
| --- | --- | --- | --- | --- | --- | --- |
| bioRxiv | new applications | 0.781 | 0.630 | 2.21 | 52.0 | 17 |
| bioRxiv | new applications + AIDS | 0.793 | 0.647 | 2.31 | 49.7 | 17 |
| bioRxiv | all funded schedules (equal weight) | 0.466 | 0.217 | 0.80 | 110.2 | 17 |
| medRxiv | new applications | 0.658 | 0.449 | 2.22 | 31.7 | 17 |
| medRxiv | new applications + AIDS | 0.688 | 0.478 | 2.36 | 30.0 | 17 |
| medRxiv | all funded schedules (equal weight) | 0.330 | 0.103 | 0.72 | 51.6 | 17 |

**Table S14.** Moving the individual surges away from the due dates (falsification of the kernel fit). The investigator kernel is moved by the stated number of weeks (positive, earlier in the cycle) before each new-application due date and fitted at each shift with one amplitude; fits with a negative amplitude invert the model and are not admissible. The last row of each server is a sinusoid with the period of the cycle and a free phase.

| Server | Kernel shift (weeks) | Weighted $R^2$ | Note |
| --- | --- | --- | --- |
| bioRxiv | -8 | – | negative amplitude |
| bioRxiv | -7 | – | negative amplitude |
| bioRxiv | -6 | – | negative amplitude |
| bioRxiv | -5 | – | negative amplitude |
| bioRxiv | -4 | – | negative amplitude |
| bioRxiv | -3 | 0.016 |  |
| bioRxiv | -2 | 0.233 |  |
| bioRxiv | -1 | 0.582 |  |
| bioRxiv | +0 | 0.630 |  |
| bioRxiv | +1 | 0.487 |  |
| bioRxiv | +2 | 0.373 |  |
| bioRxiv | +3 | 0.249 |  |
| bioRxiv | +4 | 0.034 |  |
| bioRxiv | +5 | – | negative amplitude |
| bioRxiv | +6 | – | negative amplitude |
| bioRxiv | +7 | – | negative amplitude |
| bioRxiv | +8 | – | negative amplitude |
| bioRxiv | free-phase sinusoid | 0.774 | three parameters |
| medRxiv | -8 | – | negative amplitude |
| medRxiv | -7 | – | negative amplitude |
| medRxiv | -6 | – | negative amplitude |
| medRxiv | -5 | – | negative amplitude |
| medRxiv | -4 | – | negative amplitude |
| medRxiv | -3 | – | negative amplitude |
| medRxiv | -2 | 0.029 |  |
| medRxiv | -1 | 0.220 |  |
| medRxiv | +0 | 0.449 |  |
| medRxiv | +1 | 0.550 |  |
| medRxiv | +2 | 0.551 |  |
| medRxiv | +3 | 0.340 |  |
| medRxiv | +4 | 0.179 |  |
| medRxiv | +5 | 0.030 |  |
| medRxiv | +6 | – | negative amplitude |
| medRxiv | +7 | – | negative amplitude |
| medRxiv | +8 | – | negative amplitude |
| medRxiv | free-phase sinusoid | 0.652 | three parameters |

**Table S15.** National drop at the R01 date, the resubmission date and a placebo date 45 days after the R01 date. Local linear discontinuity on US vs non-US daily counts with US-specific slopes on each side (as table S22); resubmission dates on the business day. Individual surges placed before new-application and resubmission due dates with separate amplitudes: bioRxiv: per funded resubmission, 0.15 of the amplitude per funded new application, weighted *R*^2^ 0.648; medRxiv: per funded resubmission, 0.02 of the amplitude per funded new application, weighted *R*^2^ 0.449.

| Server | Date | Cycles | Bandwidth (days) | Drop (SE) |
| --- | --- | --- | --- | --- |
| bioRxiv | R01 date | Pooled | 7 | 0.100 (0.048) |
| bioRxiv | R01 date | Pooled | 10 | 0.150 (0.049) |
| bioRxiv | Resubmission date | Pooled | 7 | 0.035 (0.052) |
| bioRxiv | Resubmission date | Pooled | 10 | 0.080 (0.043) |
| bioRxiv | Resubmission date | March | 7 | 0.003 (0.063) |
| bioRxiv | Resubmission date | March | 10 | 0.019 (0.070) |
| bioRxiv | Resubmission date | July | 7 | 0.092 (0.126) |
| bioRxiv | Resubmission date | July | 10 | 0.129 (0.090) |
| bioRxiv | Resubmission date | November | 7 | 0.037 (0.080) |
| bioRxiv | Resubmission date | November | 10 | 0.120 (0.052) |
| bioRxiv | Placebo, 45 days after r01 date | Pooled | 7 | 0.018 (0.052) |
| bioRxiv | Placebo, 45 days after r01 date | Pooled | 10 | 0.060 (0.045) |
| medRxiv | R01 date | Pooled | 7 | -0.059 (0.145) |
| medRxiv | R01 date | Pooled | 10 | 0.057 (0.093) |
| medRxiv | Resubmission date | Pooled | 7 | -0.095 (0.089) |
| medRxiv | Resubmission date | Pooled | 10 | -0.076 (0.078) |
| medRxiv | Resubmission date | March | 7 | -0.112 (0.104) |
| medRxiv | Resubmission date | March | 10 | -0.006 (0.070) |
| medRxiv | Resubmission date | July | 7 | -0.242 (0.188) |
| medRxiv | Resubmission date | July | 10 | -0.184 (0.156) |
| medRxiv | Resubmission date | November | 7 | -0.112 (0.219) |
| medRxiv | Resubmission date | November | 10 | -0.146 (0.187) |
| medRxiv | Placebo, 45 days after r01 date | Pooled | 7 | -0.126 (0.119) |
| medRxiv | Placebo, 45 days after r01 date | Pooled | 10 | -0.071 (0.091) |

**Table S16.** bioRxiv subject categories ordered by NIH dependence. Change in US share in the R01 and F windows (pp, cycle fixed effects) and the US *×* R01-window coefficient on log daily counts. Slopes on dependence (share-based): Weighted, all categories (heteroskedasticity-robust se): slope 3.14 (SE 1.13). Leave-one-out minimum (plant biology dropped): slope 2.38 (SE 1.10). Leave-one-out maximum (neuroscience dropped): slope 4.23 (SE 0.80). First half of the period: slope 3.87 (SE 0.77). Second half of the period: slope 1.81 (SE 1.60). Inverse-variance means: dependence below 0.6 (5 categories) 1.19 (0.46) pp, 0.6 or more (16 categories) 2.38 (0.23) pp, difference 1.19 (0.51) pp with categories treated as independent; Spearman correlation with dependence within the second group 0.30 (*P* = 0.26).

| Category | First versions | NIH dependence | R01 window, pp | F window, pp | R01, log counts |
| --- | --- | --- | --- | --- | --- |
| Ecology | 14,616 | 0.19 | 0.70 (0.88) | -1.14 (0.91) | 0.032 (0.045) |
| Plant biology | 12,392 | 0.38 | 0.22 (0.85) | -0.30 (0.87) | 0.013 (0.048) |
| Evolutionary biology | 15,079 | 0.47 | 1.48 (0.84) | -1.23 (0.94) | 0.061 (0.038) |
| Animal behavior and cognition | 5,374 | 0.52 | 4.05 (1.44) | -2.41 (1.50) | 0.177 (0.058) |
| Synthetic biology | 3,329 | 0.55 | 1.90 (1.99) | -0.51 (2.02) | 0.059 (0.080) |
| Microbiology | 29,987 | 0.69 | 3.10 (0.65) | -1.80 (0.71) | 0.126 (0.032) |
| Bioengineering | 11,446 | 0.71 | 2.01 (1.12) | -1.70 (1.16) | 0.077 (0.037) |
| Bioinformatics | 28,160 | 0.72 | 1.35 (0.66) | 0.05 (0.67) | 0.056 (0.024) |
| Systems biology | 6,750 | 0.73 | 1.55 (1.37) | -3.03 (1.40) | 0.053 (0.054) |
| Genomics | 15,390 | 0.74 | 3.42 (0.93) | -1.60 (0.96) | 0.136 (0.040) |
| Biophysics | 15,602 | 0.74 | 2.67 (0.93) | -2.32 (0.99) | 0.106 (0.041) |
| Pharmacology and toxicology | 3,868 | 0.75 | 3.32 (1.86) | -5.76 (1.75) | 0.129 (0.086) |
| Physiology | 6,268 | 0.77 | 3.45 (1.50) | -2.85 (1.46) | 0.139 (0.060) |
| Genetics | 11,036 | 0.79 | 3.97 (1.06) | -4.32 (1.13) | 0.155 (0.050) |
| Neuroscience | 62,371 | 0.79 | 0.95 (0.51) | -1.64 (0.48) | 0.037 (0.023) |
| Developmental biology | 10,159 | 0.80 | 2.67 (1.07) | -2.04 (1.10) | 0.107 (0.044) |
| Biochemistry | 13,959 | 0.81 | 3.51 (0.97) | -2.94 (1.02) | 0.145 (0.054) |
| Molecular biology | 13,348 | 0.82 | 2.47 (0.95) | -4.08 (0.87) | 0.107 (0.041) |
| Cancer biology | 14,925 | 0.82 | 2.78 (1.02) | -1.61 (1.08) | 0.106 (0.040) |
| Cell biology | 19,882 | 0.83 | 2.70 (0.91) | -2.71 (0.84) | 0.113 (0.043) |
| Immunology | 12,465 | 0.84 | 3.76 (1.16) | -1.65 (1.21) | 0.149 (0.050) |

**Table S17.** Preprints posted 0–13 days before the investigator’s own due date compared with the same investigator’s other preprints (investigator and year-month fixed effects, SE clustered by investigator).

| Outcome | Difference (SE) | Comparison mean | N | Pre-deadline |
| --- | --- | --- | --- | --- |
| Results first \|log ratio\| | −0.490 (0.250) | 1.411 | 5,919 | 63 |
| promotional adjectives per 1,000 words (without “significant”) | −0.776 (0.348) | 10.512 | 89,074 | 763 |
| log10 sample size | 0.130 (0.121) | 2.747 | 9,801 | 84 |
| share $P < .05$ | 0.103 (0.102) | 0.878 | 3,715 | 38 |
| versions posted | −0.003 (0.027) | 1.355 | 89,075 | 763 |
| in a journal within 24 months | −0.032 (0.023) | 0.598 | 62,979 | 596 |
| log days to journal (published only) | 0.057 (0.051) | 5.234 | 47,024 | 405 |
| log(1+citations) | −0.093 (0.051) | 1.874 | 89,075 | 763 |
| similarity to the upcoming award | 0.035 (0.008) | 0.162 | 11,321 | 763 |
| similarity to PI’s other awards | −0.000 (0.004) | 0.144 | 10,733 | 712 |
| upcoming minus other awards | 0.032 (0.008) | 0.017 | 10,733 | 712 |
| similarity to the upcoming award, adjusting for log days to due date | 0.029 (0.010) | − | 11,321 | 763 |
| upcoming minus other awards, adjusting for log days to due date | 0.029 (0.011) | − | 10,733 | 712 |
| promotional adjectives per 1,000 words (without “significant”), adjusting for log days to due date | −0.905 (0.680) | − | 11,321 | 763 |
| Results \|log ratio\|, adjusting for log days to due date | 1.239 (0.575) | − | 768 | 63 |
| similarity to the upcoming award, within 90 days of due date | 0.028 (0.013) | − | 3,451 | 763 |
| upcoming minus other awards, within 90 days of due date | 0.032 (0.013) | − | 3,259 | 712 |
| promotional adjectives per 1,000 words (without “significant”), within 90 days of due date | −1.525 (0.826) | − | 3,451 | 763 |
| Results \|log ratio\|, within 90 days of due date | 0.194 (0.299) | − | 259 | 63 |
| similarity to the upcoming award, within 45 days of due date | 0.021 (0.020) | − | 1,990 | 763 |
| upcoming minus other awards, within 45 days of due date | 0.031 (0.020) | − | 1,870 | 712 |
| promotional adjectives per 1,000 words (without “significant”), within 45 days of due date | −1.283 (1.106) | − | 1,990 | 763 |
| Results \|log ratio\|, within 45 days of due date | 0.194 (0.280) | − | 167 | 63 |

**Table S18.** Citations, publication and promotional language of pre-deadline preprints. Citation percentile: citations to a first version and its journal version in the calendar year of posting and the two following years, ranked within server, subject category and posting month (0 to 100; first versions posted through 2023). Upper rows: linked investigators’ preprints posted 0–13 days before their own due date against the same investigator’s other preprints (investigator and year-month fixed effects, SE clustered by investigator); similarity is split at the median similarity to the forthcoming award among all compared preprints; publication is a journal version linked by the server by June 2026; promotional adjectives use the 139-word list of Millar and colleagues, unchanged and without “significant”. Last rows: US *×* R01 window on all country-coded first versions (category-by-month fixed effects, SE clustered by cycle), divided by the share of US window posts the deadline adds (0.09 bioRxiv, 0.11 medRxiv).

| Comparison | Estimate (SE) | N | Pre-deadline preprints |
| --- | --- | --- | --- |
| Citation percentile, all pre-deadline preprints | −2.65 (1.41) | 56,780 | 556 |
| Top 10% of field and month (share) | −0.02 (0.02) | 56,780 | 556 |
| Citation percentile, similarity to the award at or below the median | −8.04 (2.68) | 8,192 | 210 |
| Difference, high minus low similarity | 8.48 (3.68) | 8,192 | 556 |
| Citation percentile, similarity to the award above the median | 0.44 (2.77) | 8,192 | 346 |
| Published by June 2026 (share) | −0.034 (0.023) | 56,780 | 556 |
| Citation percentile, published preprints | −0.53 (1.48) | 36,925 | 344 |
| Citation percentile, not published preprints | −4.32 (2.27) | 19,855 | 212 |
| Promotional adjectives per 1,000 words, full published list | −0.83 (0.36) | 89,074 | 763 |
| Promotional adjectives per 1,000 words, list without ‘significant’ | −0.78 (0.35) | 89,074 | 763 |
| bioRxiv: US × R01 window, all first versions | −0.42 (0.33) | 202,595 | — |
| bioRxiv: implied difference for the posts the deadline adds | −4.67 (3.73) | — | — |
| medRxiv: US × R01 window, all first versions | 1.03 (0.73) | 26,812 | — |
| medRxiv: implied difference for the posts the deadline adds | 9.32 (6.63) | — | — |

**Table S19.** Total preprints per investigator-cycle around applications (Poisson, investigator and calendar-cycle fixed effects, cycles with no preprints kept). Output is higher in the cycles around an application.

| Design | Cycle before | Own cycle | Cycle after | Sum |
| --- | --- | --- | --- | --- |
| Funded new applications | 0.072 (0.020) | 0.142 (0.020) | 0.052 (0.019) | 0.266 (0.038) |
| Failed first submission | 0.062 (0.021) | −0.005 (0.024) | 0.047 (0.020) | 0.104 (0.026) |
| Funded renewals | 0.180 (0.027) | 0.233 (0.027) | 0.064 (0.027) | 0.476 (0.048) |
| Renewal clock, all R01s | 0.073 (0.016) | 0.130 (0.016) | 0.129 (0.016) | 0.332 (0.030) |

**Table S20.** US first versions in R01 windows above, and in F windows below, the number expected from non-US counts and the US-to-non-US ratio outside the R01 and F windows in the same year (2026: January–June).

| Year | US first versions | R01-window excess | F-window deficit | Excess, % of US |
| --- | --- | --- | --- | --- |
| 2017 | 4,997 | 45 | -79 | 0.9 |
| 2018 | 8,377 | -30 | -151 | -0.4 |
| 2019 | 12,099 | 182 | -303 | 1.5 |
| 2020 | 15,856 | 61 | -385 | 0.4 |
| 2021 | 15,555 | 324 | -140 | 2.1 |
| 2022 | 16,086 | 369 | -137 | 2.3 |
| 2023 | 18,699 | 483 | -208 | 2.6 |
| 2024 | 21,257 | 559 | -227 | 2.6 |
| 2025 | 24,616 | 429 | -376 | 1.7 |
| 2026 | 13,552 | 535 | -212 | 3.9 |

**Table S21.**
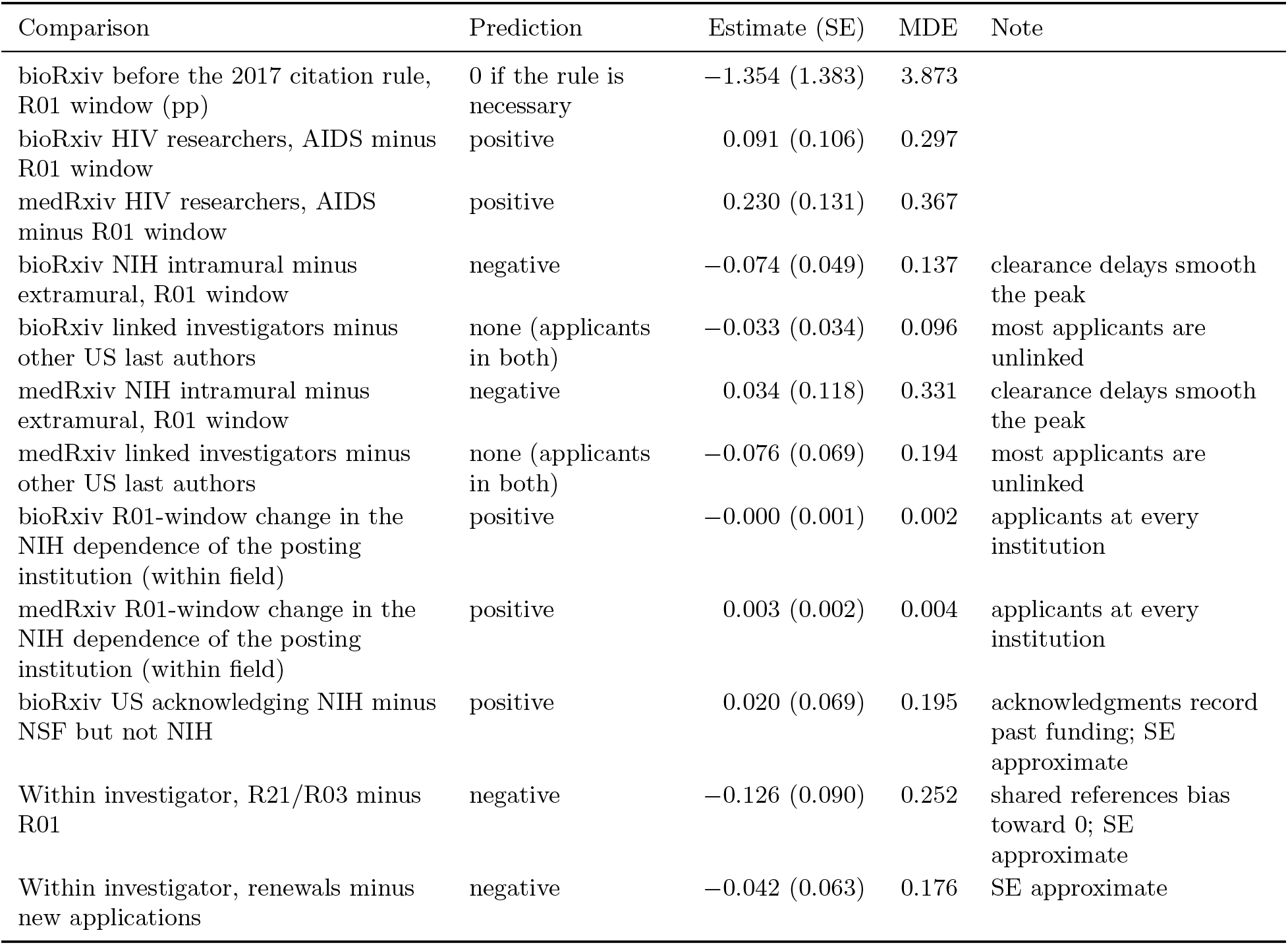
Comparisons that could count against the account. Prediction: the sign expected if the NIH calendar times the rhythm through applicants. MDE: minimum detectable effect at 80% power and a two-sided 5% level (2.8 SE). Estimates on the log scale unless stated.

| Comparison | Prediction | Estimate (SE) | MDE | Note |
| --- | --- | --- | --- | --- |
| bioRxiv before the 2017 citation rule, R01 window (pp) | 0 if the rule is necessary | -1.354 (1.383) | 3.873 |  |
| bioRxiv HIV researchers, AIDS minus R01 window | positive | 0.091 (0.106) | 0.297 |  |
| medRxiv HIV researchers, AIDS minus R01 window | positive | 0.230 (0.131) | 0.367 |  |
| bioRxiv NIH intramural minus extramural, R01 window | negative | -0.074 (0.049) | 0.137 | clearance delays smooth the peak |
| bioRxiv linked investigators minus other US last authors | none (applicants in both) | -0.033 (0.034) | 0.096 | most applicants are unlinked |
| medRxiv NIH intramural minus extramural, R01 window | negative | 0.034 (0.118) | 0.331 | clearance delays smooth the peak |
| medRxiv linked investigators minus other US last authors | none (applicants in both) | -0.076 (0.069) | 0.194 | most applicants are unlinked |
| bioRxiv R01-window change in the NIH dependence of the posting institution (within field) | positive | -0.000 (0.001) | 0.002 | applicants at every institution |
| medRxiv R01-window change in the NIH dependence of the posting institution (within field) | positive | 0.003 (0.002) | 0.004 | applicants at every institution |
| bioRxiv US acknowledging NIH minus NSF but not NIH | positive | 0.020 (0.069) | 0.195 | acknowledgments record past funding; SE approximate |
| Within investigator, R21/R03 minus R01 | negative | -0.126 (0.090) | 0.252 | shared references bias toward 0; SE approximate |
| Within investigator, renewals minus new applications | negative | -0.042 (0.063) | 0.176 | SE approximate |

**Table S22.** Discontinuity at the R01 due date. Poisson day counts of US and non-US first versions (day, group-by-cycle and group-by-weekday fixed effects, SE clustered by cycle) within the bandwidth of a due date, with a separate US linear trend on each side; the drop is the fall in the US log ratio at the due date. Business-day dates move a due date that falls on a weekend to the following Monday. Rank *P* compares the drop with the same estimate at due dates shifted by 14 to 60 days (94 shifts).

| Server | Due dates | Bandwidth (days) | Drop (SE) | Rank $P$ |
| --- | --- | --- | --- | --- |
| bioRxiv | Nominal | 7 | 0.106 (0.048) | — |
| bioRxiv | Nominal | 14 | 0.121 (0.037) | 0.042 |
| bioRxiv | Nominal | 21 | 0.107 (0.030) | — |
| bioRxiv | Business day | 7 | 0.100 (0.048) | — |
| bioRxiv | Business day | 14 | 0.124 (0.035) | — |
| bioRxiv | Business day | 21 | 0.112 (0.030) | — |
| medRxiv | Nominal | 7 | −0.105 (0.130) | — |
| medRxiv | Nominal | 14 | 0.045 (0.081) | 0.305 |
| medRxiv | Nominal | 21 | 0.103 (0.081) | — |
| medRxiv | Business day | 7 | −0.059 (0.145) | — |
| medRxiv | Business day | 14 | 0.106 (0.083) | — |
| medRxiv | Business day | 21 | 0.142 (0.083) | — |

**Table S23.** Citation cost of deadline crowding (log(1 + citations), field-by-year and calendar-week fixed effects, SE clustered by week) and acknowledgment of the forthcoming award (linear probability, investigator and year-month fixed effects).

| Sample | Term | Estimate (SE) | N |
| --- | --- | --- | --- |
| all authors | R01 window x field NIH share | 0.014 (0.024) | 262,098 |
| all authors | log same-field same-week preprints | −0.029 (0.013) | 262,098 |
| non-US authors | R01 window x field NIH share | 0.019 (0.030) | 154,575 |
| non-US authors | log same-field same-week preprints | −0.009 (0.017) | 154,575 |
| US authors | R01 window x field NIH share | −0.007 (0.035) | 107,523 |
| US authors | log same-field same-week preprints | −0.064 (0.021) | 107,523 |
| Award acknowledgment | preprint acknowledges the award (adjusting for log days to due date) | −0.006 (0.023) | 11,321 |
| Award acknowledgment | preprint acknowledges the award (within 90 days of due date) | −0.029 (0.027) | 3,451 |
| Award acknowledgment | preprint acknowledges the award (journal version exists, adjusting for log days) | −0.016 (0.036) | 6,733 |
| Award acknowledgment | journal version acknowledges the award (journal version exists, adjusting for log days) | 0.060 (0.044) | 6,733 |
| Award acknowledgment | either acknowledges the award (adjusting for log days to due date) | 0.027 (0.029) | 11,321 |
| Award acknowledgment | either acknowledges the award (within 90 days of due date) | 0.005 (0.033) | 3,451 |
| Award acknowledgment | either acknowledges the award (journal version exists, adjusting for log days) | 0.063 (0.044) | 6,733 |

**Table S24.** Consistency checks on daily counts (Poisson). Model A: per-group rate ratio, cycle and weekday fixed effects. Model B: relative to the reference group on the same day, with day, group-by-cycle and group-by-weekday fixed effects. SE clustered by cycle.

| Comparison | Model | Group or term | Window | Estimate (SE) |
| --- | --- | --- | --- | --- |
| bioRxiv HIV (all days) | A | US-HIV | R01 | 0.076 (0.053) |
| bioRxiv HIV (all days) | A | US-HIV | AIDS | 0.089 (0.049) |
| bioRxiv HIV (all days) | A | US-HIV | F | −0.047 (0.038) |
| bioRxiv HIV (all days) | A | US-other | R01 | 0.141 (0.018) |
| bioRxiv HIV (all days) | A | US-other | AIDS | −0.101 (0.016) |
| bioRxiv HIV (all days) | A | US-other | F | −0.060 (0.013) |
| bioRxiv HIV (all days) | A | nonUS-HIV | R01 | −0.029 (0.082) |
| bioRxiv HIV (all days) | A | nonUS-HIV | AIDS | −0.024 (0.060) |
| bioRxiv HIV (all days) | A | nonUS-HIV | F | 0.074 (0.065) |
| bioRxiv HIV (all days) | A | nonUS-other | R01 | 0.054 (0.018) |
| bioRxiv HIV (all days) | A | nonUS-other | AIDS | −0.074 (0.012) |
| bioRxiv HIV (all days) | A | nonUS-other | F | 0.021 (0.014) |
| bioRxiv HIV (all days) | B | US | R01 | 0.085 (0.014) |
| bioRxiv HIV (all days) | B | HIV | R01 | −0.088 (0.077) |
| bioRxiv HIV (all days) | B | US × HIV | R01 | 0.020 (0.091) |
| bioRxiv HIV (all days) | B | US | AIDS | −0.026 (0.013) |
| bioRxiv HIV (all days) | B | HIV | AIDS | 0.050 (0.058) |
| bioRxiv HIV (all days) | B | US × HIV | AIDS | 0.140 (0.084) |
| bioRxiv HIV (all days) | B | US | F | −0.081 (0.011) |
| bioRxiv HIV (all days) | B | HIV | F | 0.058 (0.063) |
| bioRxiv HIV (all days) | B | US × HIV | F | −0.042 (0.077) |
| bioRxiv HIV (conferences removed) | B | US | R01 | 0.079 (0.015) |
| bioRxiv HIV (conferences removed) | B | HIV | R01 | −0.073 (0.076) |
| bioRxiv HIV (conferences removed) | B | US × HIV | R01 | −0.024 (0.089) |
| bioRxiv HIV (conferences removed) | B | US | AIDS | −0.039 (0.013) |
| bioRxiv HIV (conferences removed) | B | HIV | AIDS | 0.045 (0.054) |
| bioRxiv HIV (conferences removed) | B | US × HIV | AIDS | 0.175 (0.077) |
| bioRxiv HIV (conferences removed) | B | US | F | −0.087 (0.012) |
| bioRxiv HIV (conferences removed) | B | HIV | F | 0.011 (0.071) |
| bioRxiv HIV (conferences removed) | B | US × HIV | F | −0.058 (0.087) |
| medRxiv HIV (all days) | A | US-HIV | R01 | −0.020 (0.077) |
| medRxiv HIV (all days) | A | US-HIV | AIDS | 0.165 (0.071) |
| medRxiv HIV (all days) | A | US-HIV | F | 0.181 (0.107) |
| medRxiv HIV (all days) | A | US-other | R01 | 0.168 (0.029) |
| medRxiv HIV (all days) | A | US-other | AIDS | −0.063 (0.037) |
| medRxiv HIV (all days) | A | US-other | F | −0.086 (0.030) |
| medRxiv HIV (all days) | A | nonUS-HIV | R01 | 0.015 (0.067) |
| medRxiv HIV (all days) | A | nonUS-HIV | AIDS | −0.022 (0.090) |
| medRxiv HIV (all days) | A | nonUS-HIV | F | 0.128 (0.094) |
| medRxiv HIV (all days) | A | nonUS-other | R01 | 0.064 (0.022) |
| medRxiv HIV (all days) | A | nonUS-other | AIDS | -0.069 (0.030) |
| medRxiv HIV (all days) | A | nonUS-other | F | 0.019 (0.018) |
| medRxiv HIV (all days) | B | US | R01 | 0.104 (0.027) |
| medRxiv HIV (all days) | B | HIV | R01 | -0.055 (0.065) |
| medRxiv HIV (all days) | B | US × HIV | R01 | -0.124 (0.096) |
| medRxiv HIV (all days) | B | US | AIDS | 0.007 (0.030) |
| medRxiv HIV (all days) | B | HIV | AIDS | 0.051 (0.080) |
| medRxiv HIV (all days) | B | US × HIV | AIDS | 0.180 (0.123) |
| medRxiv HIV (all days) | B | US | F | -0.105 (0.026) |
| medRxiv HIV (all days) | B | HIV | F | 0.111 (0.093) |
| medRxiv HIV (all days) | B | US × HIV | F | 0.146 (0.126) |
| bioRxiv NIH intramural | A | US-extramural | R01 | 0.152 (0.018) |
| bioRxiv NIH intramural | A | US-extramural | AIDS | -0.105 (0.014) |
| bioRxiv NIH intramural | A | US-extramural | F | -0.066 (0.011) |
| bioRxiv NIH intramural | A | US-intramural | R01 | 0.077 (0.052) |
| bioRxiv NIH intramural | A | US-intramural | AIDS | -0.066 (0.045) |
| bioRxiv NIH intramural | A | US-intramural | F | -0.045 (0.046) |
| bioRxiv NIH intramural | A | nonUS | R01 | 0.056 (0.016) |
| bioRxiv NIH intramural | A | nonUS | AIDS | -0.070 (0.012) |
| bioRxiv NIH intramural | A | nonUS | F | 0.013 (0.015) |
| bioRxiv NIH intramural | B | US-intramural | R01 | -0.074 (0.049) |
| bioRxiv NIH intramural | B | nonUS | R01 | -0.093 (0.014) |
| bioRxiv NIH intramural | B | US-intramural | AIDS | 0.040 (0.046) |
| bioRxiv NIH intramural | B | nonUS | AIDS | 0.034 (0.013) |
| bioRxiv NIH intramural | B | US-intramural | F | 0.019 (0.048) |
| bioRxiv NIH intramural | B | nonUS | F | 0.078 (0.012) |
| medRxiv NIH intramural | A | US-extramural | R01 | 0.169 (0.023) |
| medRxiv NIH intramural | A | US-extramural | AIDS | -0.055 (0.031) |
| medRxiv NIH intramural | A | US-extramural | F | -0.083 (0.029) |
| medRxiv NIH intramural | A | US-intramural | R01 | 0.202 (0.129) |
| medRxiv NIH intramural | A | US-intramural | AIDS | 0.079 (0.112) |
| medRxiv NIH intramural | A | US-intramural | F | -0.160 (0.089) |
| medRxiv NIH intramural | A | nonUS | R01 | 0.053 (0.019) |
| medRxiv NIH intramural | A | nonUS | AIDS | -0.053 (0.030) |
| medRxiv NIH intramural | A | nonUS | F | 0.024 (0.016) |
| medRxiv NIH intramural | B | US-intramural | R01 | 0.034 (0.118) |
| medRxiv NIH intramural | B | nonUS | R01 | -0.116 (0.023) |
| medRxiv NIH intramural | B | US-intramural | AIDS | 0.126 (0.102) |
| medRxiv NIH intramural | B | nonUS | AIDS | 0.002 (0.023) |
| medRxiv NIH intramural | B | US-intramural | F | -0.069 (0.085) |
| medRxiv NIH intramural | B | nonUS | F | 0.108 (0.024) |
| bioRxiv linked investigators | A | US-linked-PI | R01 | 0.134 (0.023) |
| bioRxiv linked investigators | A | US-linked-PI | AIDS | -0.119 (0.020) |
| bioRxiv linked investigators | A | US-linked-PI | F | -0.086 (0.016) |
| bioRxiv linked investigators | A | US-other | R01 | 0.173 (0.026) |
| bioRxiv linked investigators | A | US-other | AIDS | -0.084 (0.018) |
| bioRxiv linked investigators | A | US-other | F | -0.037 (0.018) |
| bioRxiv linked investigators | A | nonUS | R01 | 0.056 (0.016) |
| bioRxiv linked investigators | A | nonUS | AIDS | -0.070 (0.012) |
| bioRxiv linked investigators | A | nonUS | F | 0.013 (0.015) |
| bioRxiv linked investigators | B | US-linked-PI | R01 | -0.033 (0.034) |
| bioRxiv linked investigators | B | nonUS | R01 | -0.111 (0.023) |
| bioRxiv linked investigators | B | US-linked-PI | AIDS | −0.033 (0.025) |
| bioRxiv linked investigators | B | nonUS | AIDS | 0.014 (0.017) |
| bioRxiv linked investigators | B | US-linked-PI | F | −0.061 (0.035) |
| bioRxiv linked investigators | B | nonUS | F | 0.043 (0.028) |
| medRxiv linked investigators | A | US-linked-PI | R01 | 0.128 (0.047) |
| medRxiv linked investigators | A | US-linked-PI | AIDS | −0.081 (0.042) |
| medRxiv linked investigators | A | US-linked-PI | F | −0.092 (0.053) |
| medRxiv linked investigators | A | US-other | R01 | 0.200 (0.034) |
| medRxiv linked investigators | A | US-other | AIDS | −0.032 (0.040) |
| medRxiv linked investigators | A | US-other | F | −0.080 (0.029) |
| medRxiv linked investigators | A | nonUS | R01 | 0.053 (0.019) |
| medRxiv linked investigators | A | nonUS | AIDS | −0.053 (0.030) |
| medRxiv linked investigators | A | nonUS | F | 0.024 (0.016) |
| medRxiv linked investigators | B | US-linked-PI | R01 | −0.076 (0.069) |
| medRxiv linked investigators | B | nonUS | R01 | −0.149 (0.040) |
| medRxiv linked investigators | B | US-linked-PI | AIDS | −0.035 (0.049) |
| medRxiv linked investigators | B | nonUS | AIDS | −0.016 (0.033) |
| medRxiv linked investigators | B | US-linked-PI | F | −0.011 (0.061) |
| medRxiv linked investigators | B | nonUS | F | 0.104 (0.030) |
| bioRxiv HIV, joint model | B | US × HIV × AIDS | − | 0.160 (0.089) |
| bioRxiv HIV, joint model | B | US × HIV × R new | − | 0.070 (0.096) |
| bioRxiv HIV, joint model | B | phase shift: US × HIV ×<br>(AIDS - R01) | − | 0.091 (0.106) |
| medRxiv HIV, joint model | B | US × HIV × AIDS | − | 0.158 (0.132) |
| medRxiv HIV, joint model | B | US × HIV × R new | − | −0.072 (0.105) |
| medRxiv HIV, joint model | B | phase shift: US × HIV ×<br>(AIDS - R01) | − | 0.230 (0.131) |

**Table S25.** The drop at the R01 due date without East Asian authors in the comparison group (CN, HK, MO, TW, KR, SG, VN) and without the Lunar New Year holiday weeks and 1–7 October, which lower posting in East Asia near the February and October due dates. Country-coded first versions; drop and rank *P* as in table S2; discontinuity with a 14-day bandwidth as in table S22.

| Server | Comparison group | Holidays | Cycles | Drop (SE) | Rank $P$ | Discontinuity (SE) |
| --- | --- | --- | --- | --- | --- | --- |
| bioRxiv | all Non-US | kept | all | 0.112 (0.030) | 0.011 | 0.130 (0.037) |
| bioRxiv | all Non-US | kept | February | 0.178 (0.039) | 0.011 | 0.109 (0.100) |
| bioRxiv | all Non-US | removed | all | 0.122 (0.036) | 0.011 | 0.157 (0.048) |
| bioRxiv | all Non-US | removed | February | 0.165 (0.043) | 0.011 | 0.159 (0.104) |
| bioRxiv | Non-US without East Asia | kept | all | 0.118 (0.032) | 0.011 | 0.137 (0.039) |
| bioRxiv | Non-US without East Asia | kept | February | 0.168 (0.039) | 0.021 | 0.101 (0.097) |
| bioRxiv | Non-US without East Asia | removed | all | 0.127 (0.037) | 0.011 | 0.163 (0.050) |
| bioRxiv | Non-US without East Asia | removed | February | 0.160 (0.043) | 0.032 | 0.149 (0.110) |
| medRxiv | all Non-US | kept | all | 0.118 (0.056) | 0.032 | 0.072 (0.088) |
| medRxiv | all Non-US | kept | February | 0.133 (0.065) | 0.084 | 0.174 (0.148) |
| medRxiv | all Non-US | removed | all | 0.124 (0.072) | 0.032 | 0.119 (0.119) |
| medRxiv | all Non-US | removed | February | 0.082 (0.103) | 0.221 | 0.334 (0.251) |
| medRxiv | Non-US without East Asia | kept | all | 0.117 (0.059) | 0.032 | 0.075 (0.089) |
| medRxiv | Non-US without East Asia | kept | February | 0.110 (0.070) | 0.147 | 0.219 (0.150) |
| medRxiv | Non-US without East Asia | removed | all | 0.131 (0.078) | 0.032 | 0.139 (0.126) |
| medRxiv | Non-US without East Asia | removed | February | 0.064 (0.109) | 0.305 | 0.380 (0.262) |

**Table S26.** Week-by-week field gradient on bioRxiv daily counts. For each 7-day bin of days to the next R01 due date, the per-field US *×* bin coefficient regressed on NIH dependence (21 fields, inverse-variance weights, HC1 SE). The intercept is the coefficient expected for a field with no NIH dependence.

| Bin | Days to R01 due date | Intercept (SE) | Slope (SE) |
| --- | --- | --- | --- |
| 0 | 0–6 | −0.027 (0.045) | 0.215 (0.079) |
| 1 | 7–13 | 0.004 (0.080) | 0.102 (0.121) |
| 2 | 14–20 | −0.045 (0.054) | 0.110 (0.081) |
| 3 | 21–27 | 0.106 (0.039) | −0.108 (0.078) |
| 4 | 28–34 | 0.020 (0.066) | −0.016 (0.091) |
| 5 | 35–41 | 0.033 (0.047) | −0.046 (0.067) |
| 6 | 42–48 | −0.141 (0.054) | 0.103 (0.074) |
| 7 | 49–55 | −0.017 (0.077) | −0.018 (0.105) |
| 8 | 56–62 | 0.010 (0.050) | −0.037 (0.073) |
| 9 | 63–69 | −0.029 (0.055) | −0.047 (0.075) |
| 10 | 70–76 | −0.066 (0.057) | −0.107 (0.085) |
| 11 | 77–83 | 0.085 (0.049) | −0.199 (0.069) |
| 12 | 84–90 | 0.052 (0.152) | −0.133 (0.200) |
| 13 | 91–97 | −0.158 (0.068) | 0.207 (0.091) |
| 14 | 98–104 | −0.024 (0.072) | 0.054 (0.106) |
| 15 | 105–111 | 0.035 (0.041) | −0.046 (0.062) |
| 16 | 112–118 | 0.045 (0.044) | −0.038 (0.067) |
| 17 | 119–125 | 0.107 (0.093) | 0.001 (0.138) |

**Table S27.**
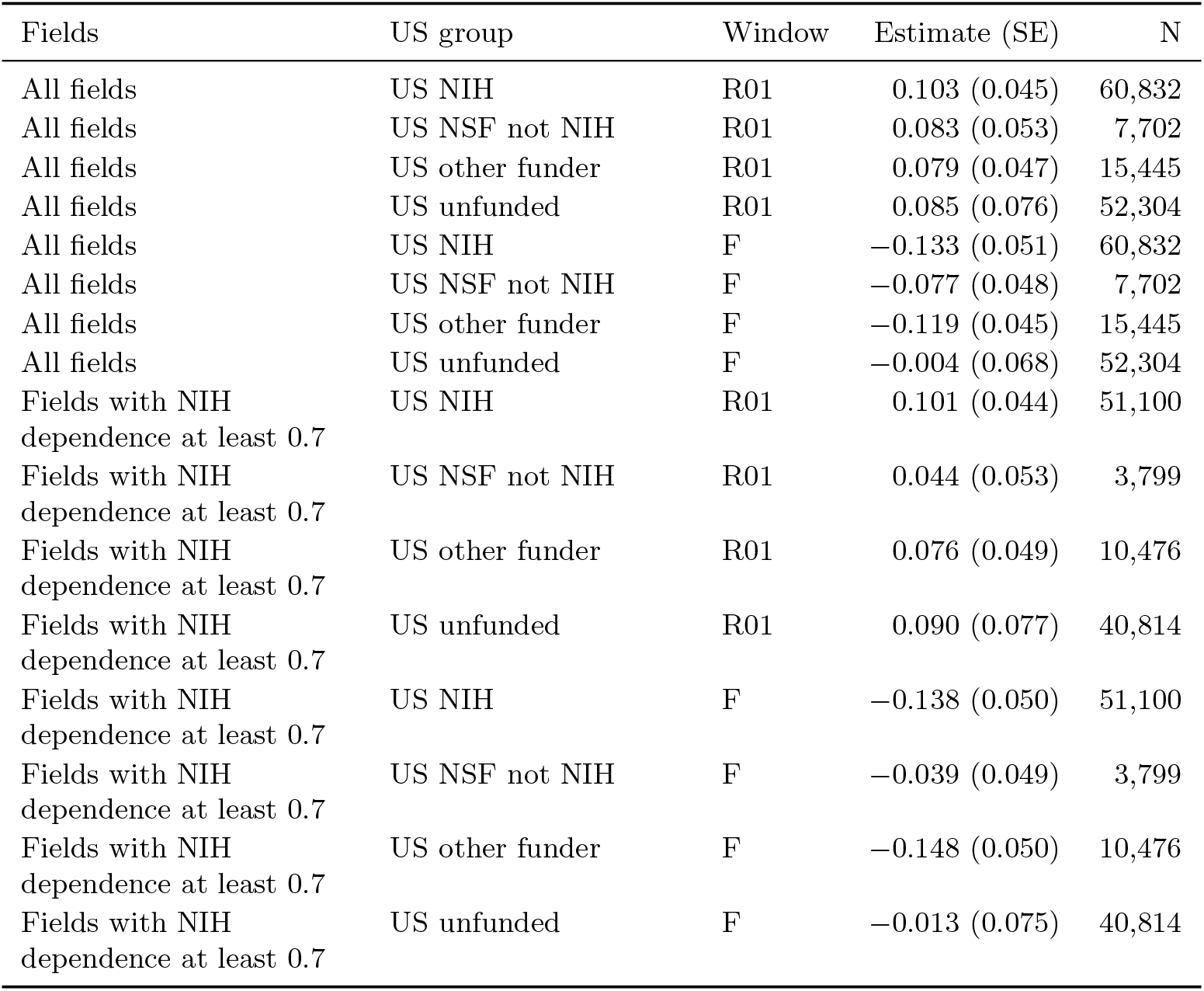
US bioRxiv first versions by OpenAlex funder group, each relative to non-US first versions on the same day (Poisson, day, group-by-cycle and group-by-weekday fixed effects). All US groups rise in the R01 window, by amounts that cannot be distinguished.

| Fields | US group | Window | Estimate (SE) | N |
| --- | --- | --- | --- | --- |
| All fields | US NIH | R01 | 0.103 (0.045) | 60,832 |
| All fields | US NSF not NIH | R01 | 0.083 (0.053) | 7,702 |
| All fields | US other funder | R01 | 0.079 (0.047) | 15,445 |
| All fields | US unfunded | R01 | 0.085 (0.076) | 52,304 |
| All fields | US NIH | F | −0.133 (0.051) | 60,832 |
| All fields | US NSF not NIH | F | −0.077 (0.048) | 7,702 |
| All fields | US other funder | F | −0.119 (0.045) | 15,445 |
| All fields | US unfunded | F | −0.004 (0.068) | 52,304 |
| Fields with NIH dependence at least 0.7 | US NIH | R01 | 0.101 (0.044) | 51,100 |
| Fields with NIH dependence at least 0.7 | US NSF not NIH | R01 | 0.044 (0.053) | 3,799 |
| Fields with NIH dependence at least 0.7 | US other funder | R01 | 0.076 (0.049) | 10,476 |
| Fields with NIH dependence at least 0.7 | US unfunded | R01 | 0.090 (0.077) | 40,814 |
| Fields with NIH dependence at least 0.7 | US NIH | F | −0.138 (0.050) | 51,100 |
| Fields with NIH dependence at least 0.7 | US NSF not NIH | F | −0.039 (0.049) | 3,799 |
| Fields with NIH dependence at least 0.7 | US other funder | F | −0.148 (0.050) | 10,476 |
| Fields with NIH dependence at least 0.7 | US unfunded | F | −0.013 (0.075) | 40,814 |

**Table S28.** From the across-investigator estimate to the within-investigator estimate (funded new applications). The Poisson model with investigator-by-cycle fixed effects estimates a log odds ratio of the window share; share models report log(1 + *b/ȳ*).

| Window (days) | Specification | Estimate (SE) |
| --- | --- | --- |
| 28 | across Poisson (investigator-by-cycle and cycle-by-window FE), all units | 0.178 (0.042) |
| 28 | across Poisson (investigator-by-cycle and cycle-by-window FE), units with at least 2 cycles | 0.156 (0.044) |
| 28 | share regression, calendar-cycle FE only, all units | 0.130 (0.034) |
| 28 | share regression, calendar-cycle FE only, units with at least 2 cycles | 0.114 (0.035) |
| 28 | share regression, investigator and calendar-cycle FE (within) | 0.087 (0.039) |
| 28 | Poisson window count, offset log N, calendar-cycle FE only | 0.129 (0.029) |
| 28 | Poisson window count, offset log N, investigator and calendar-cycle FE (within) | 0.084 (0.036) |
| 14 | across Poisson (investigator-by-cycle and cycle-by-window FE), all units | 0.213 (0.050) |
| 14 | across Poisson (investigator-by-cycle and cycle-by-window FE), units with at least 2 cycles | 0.181 (0.053) |
| 14 | share regression, calendar-cycle FE only, all units | 0.185 (0.051) |
| 14 | share regression, calendar-cycle FE only, units with at least 2 cycles | 0.159 (0.053) |
| 14 | share regression, investigator and calendar-cycle FE (within) | 0.115 (0.057) |
| 14 | Poisson window count, offset log N, calendar-cycle FE only | 0.181 (0.042) |
| 14 | Poisson window count, offset log N, investigator and calendar-cycle FE (within) | 0.095 (0.050) |

**Table S29.**
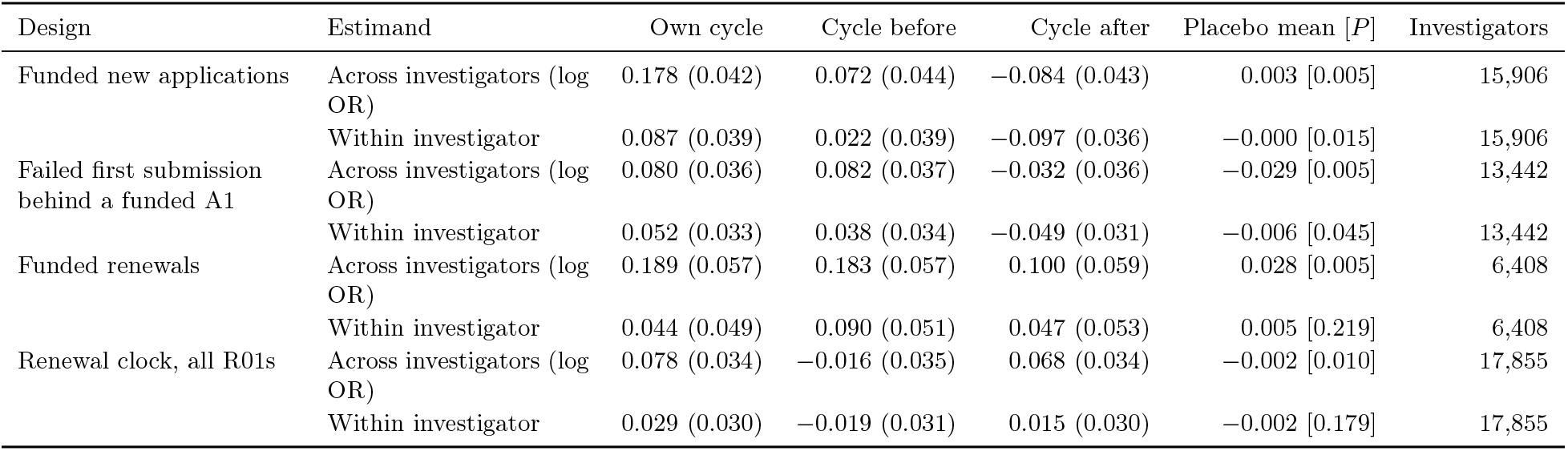
Investigator-level estimates for four designs. Own application cycle, the cycle before and the cycle after (28-day window). Placebo: mean of 200 estimates with application cycles removed and placebo cycles drawn among the remaining cycles [one-sided *P* of the observed estimate].

| Design | Estimand | Own cycle | Cycle before | Cycle after | Placebo mean [$P$] | Investigators |
| --- | --- | --- | --- | --- | --- | --- |
| Funded new applications | Across investigators (log OR) | 0.178 (0.042) | 0.072 (0.044) | -0.084 (0.043) | 0.003 [0.005] | 15,906 |
|  | Within investigator | 0.087 (0.039) | 0.022 (0.039) | -0.097 (0.036) | -0.000 [0.015] | 15,906 |
| Failed first submission behind a funded A1 | Across investigators (log OR) | 0.080 (0.036) | 0.082 (0.037) | -0.032 (0.036) | -0.029 [0.005] | 13,442 |
|  | Within investigator | 0.052 (0.033) | 0.038 (0.034) | -0.049 (0.031) | -0.006 [0.045] | 13,442 |
| Funded renewals | Across investigators (log OR) | 0.189 (0.057) | 0.183 (0.057) | 0.100 (0.059) | 0.028 [0.005] | 6,408 |
|  | Within investigator | 0.044 (0.049) | 0.090 (0.051) | 0.047 (0.053) | 0.005 [0.219] | 6,408 |
| Renewal clock, all R01s | Across investigators (log OR) | 0.078 (0.034) | -0.016 (0.035) | 0.068 (0.034) | -0.002 [0.010] | 17,855 |
|  | Within investigator | 0.029 (0.030) | -0.019 (0.031) | 0.015 (0.030) | -0.002 [0.179] | 17,855 |

**Table S30.** Simulation on the observed panel (100 replicates). Investigator-cycle totals fixed; window counts binomial with the calendar-cycle share plus investigator noise (SD 0.5 on the log-odds scale) and a known log-odds effect *β* on application cycles. Each estimator is evaluated against the truth on its own scale.

| $\beta$ | Estimator | Truth | Mean estimate | SD | Mean SE | Coverage |
| --- | --- | --- | --- | --- | --- | --- |
| 0.00 | across investigators (log odds) | 0.000 | -0.002 | 0.040 | 0.040 | 0.95 |
| 0.00 | within investigator (share scale) | 0.000 | 0.000 | 0.034 | 0.034 | 0.95 |
| 0.18 | across investigators (log odds) | 0.180 | 0.171 | 0.040 | 0.039 | 0.93 |
| 0.18 | within investigator (share scale) | 0.132 | 0.125 | 0.032 | 0.035 | 0.96 |

**Table S31.** Within-investigator estimates with the cycles immediately before and after each application removed from the reference.

| Design | Reference cycles | Estimate (SE) | Investigators |
| --- | --- | --- | --- |
| Funded new applications | all other cycles | 0.087 (0.039) | 15,906 |
| Funded new applications | excluding cycles adjacent to an application | 0.079 (0.040) | 15,682 |
| Funded renewals | all other cycles | 0.044 (0.049) | 6,408 |
| Funded renewals | excluding cycles adjacent to an application | 0.044 (0.051) | 6,252 |
| Failed first submission | all other cycles | 0.052 (0.033) | 13,442 |
| Failed first submission | excluding cycles adjacent to an application | 0.049 (0.034) | 13,195 |
| Renewal clock, all R01s | all other cycles | 0.029 (0.030) | 17,855 |
| Renewal clock, all R01s | excluding cycles adjacent to an application | 0.056 (0.031) | 17,065 |

**Table S32.** Funded new applications by activity code. R01 applications are reviewed on preliminary data; R21 and R03 applications do not require them. Investigators who hold both kinds of award keep the cycles of one kind in the reference of the other, which biases each estimate toward zero. Placebo: 100 draws among non-application cycles.

| Activity | Estimand | Estimate (SE) | Placebo 97.5% | Placebo $P$ | Investigators | Pre-deadline preprints |
| --- | --- | --- | --- | --- | --- | --- |
| R01 | Across investigators | 0.161 (0.053) | 0.066 | 0.010 | 11,221 | 626 |
| R01 | Within investigator | 0.095 (0.050) | 0.088 | 0.040 | 11,221 | 626 |
| R21/R03 | Across investigators | 0.105 (0.081) | 0.192 | 0.257 | 5,509 | 234 |
| R21/R03 | Within investigator | -0.031 (0.075) | 0.131 | 0.545 | 5,509 | 234 |

**Table S33.** Linking investigators to OpenAlex authors. Only anchored links (an author named on a paper acknowledging the investigator’s own award) were used.

|  | Value |
| --- | --- |
| Anchored links (investigators) | 98,834 (67,184) |
| Name-and-institution links (investigators) | 24,696 (21,671) |
| Name-and-institution links confirmed by anchor, all | 0.88 |
| Confirmed, stratified sample of 200 | 0.74 |
| Confirmed: uncommon surname, one OpenAlex ID(s) | 0.98 (n = 12,594) |
| Confirmed: uncommon surname, several OpenAlex ID(s) | 0.65 (n = 2,642) |
| Confirmed: common surname, one OpenAlex ID(s) | 0.89 (n = 5,458) |
| Confirmed: common surname, several OpenAlex ID(s) | 0.54 (n = 2,392) |

**Table S34.** Other countries and funders. Country shares are among all country-coded first versions, so the US rows differ slightly from table S3, which compares US with all non-US authors; SE clustered by week. Canada shares the North American academic year and may apply for R01 grants but not NIH fellowships. NSFC and KAKENHI: share of Chinese or Japanese corresponding authors among non-US first versions in the 28 days before the funder’s due date and at placebo dates.

| Server | Country | Window | Change, pp (SE) | Relative, % |
| --- | --- | --- | --- | --- |
| bioRxiv | US | R01 | 2.24 (0.28) | 5.6 |
| bioRxiv | US | F | -1.84 (0.31) | -4.6 |
| bioRxiv | CA | R01 | 0.10 (0.09) | 2.6 |
| bioRxiv | CA | F | -0.26 (0.08) | -6.7 |
| medRxiv | US | R01 | 2.66 (0.52) | 7.7 |
| medRxiv | US | F | -2.43 (0.54) | -7.0 |
| medRxiv | CA | R01 | 0.52 (0.23) | 12.3 |
| medRxiv | CA | F | -0.24 (0.19) | -5.8 |
| bioRxiv | CN (NSFC) | own due date | 0.63 (0.37) | 6.7 |
| bioRxiv | CN (NSFC) | placebo -182 d | -0.72 (0.30) | -7.8 |
| bioRxiv | CN (NSFC) | placebo -91 d | 0.03 (0.31) | 0.3 |
| bioRxiv | CN (NSFC) | placebo +91 d | 0.43 (0.30) | 4.6 |
| bioRxiv | CN (NSFC) | placebo +182 d | -0.55 (0.28) | -5.9 |
| bioRxiv | JP (KAKENHI) | own due date | 0.50 (0.28) | 8.9 |
| bioRxiv | JP (KAKENHI) | placebo -182 d | -0.00 (0.27) | -0.0 |
| bioRxiv | JP (KAKENHI) | placebo -91 d | 0.20 (0.34) | 3.6 |
| bioRxiv | JP (KAKENHI) | placebo +91 d | -0.69 (0.32) | -12.4 |
| bioRxiv | JP (KAKENHI) | placebo +182 d | -0.12 (0.25) | -2.2 |
| medRxiv | CN (NSFC) | own due date | 0.87 (0.63) | 11.3 |
| medRxiv | CN (NSFC) | placebo -182 d | 0.11 (0.49) | 1.4 |
| medRxiv | CN (NSFC) | placebo -91 d | -0.63 (0.48) | -8.1 |
| medRxiv | CN (NSFC) | placebo +91 d | -0.09 (0.56) | -1.2 |
| medRxiv | CN (NSFC) | placebo +182 d | 0.01 (0.46) | 0.2 |
| medRxiv | JP (KAKENHI) | own due date | 0.15 (0.39) | 4.0 |
| medRxiv | JP (KAKENHI) | placebo -182 d | -0.09 (0.45) | -2.4 |
| medRxiv | JP (KAKENHI) | placebo -91 d | 0.58 (0.49) | 15.3 |
| medRxiv | JP (KAKENHI) | placebo +91 d | -0.13 (0.45) | -3.3 |
| medRxiv | JP (KAKENHI) | placebo +182 d | -0.38 (0.43) | -9.8 |

**Table S35.** Before and after NIH allowed applications to cite preprints (NOT-OD-17-050, 24 March 2017), bioRxiv. Country assigned in both periods from the ROR match of the corresponding institution.

| Fields | Period | Measure | Window | Estimate (SE) | N |
| --- | --- | --- | --- | --- | --- |
| All categories | pre | US share (pp) | R01 | -1.35 (1.38) | 8,290 |
| All categories | pre | US share (pp) | F | -0.23 (2.23) | 8,290 |
| All categories | pre | US share (pp) | AIDS | -0.47 (1.77) | 8,290 |
| All categories | pre | US share (pp) | Resubmission | 1.83 (1.42) | 8,290 |
| All categories | pre | log(US/nonUS) counts | R01 | -0.055 (0.060) | 8,290 |
| All categories | pre | log(US/nonUS) counts | F | -0.017 (0.090) | 8,290 |
| All categories | post | US share (pp) | R01 | 1.31 (0.91) | 54,944 |
| All categories | post | US share (pp) | F | -2.31 (0.64) | 54,944 |
| All categories | post | US share (pp) | AIDS | -1.07 (0.37) | 54,944 |
| All categories | post | US share (pp) | Resubmission | 2.00 (0.65) | 54,944 |
| All categories | post | log(US/nonUS) counts | R01 | 0.058 (0.042) | 54,944 |
| All categories | post | log(US/nonUS) counts | F | -0.104 (0.029) | 54,944 |
| All categories | difference post - pre | US share (pp) | R01 | 2.73 (1.61) | 63,234 |
| All categories | difference post - pre | US share (pp) | F | -1.63 (2.41) | 63,234 |
| High nih dependence | pre | US share (pp) | R01 | -0.30 (1.43) | 5,630 |
| High nih dependence | pre | US share (pp) | F | -0.47 (2.51) | 5,630 |
| High nih dependence | post | US share (pp) | R01 | 1.61 (0.89) | 39,436 |
| High nih dependence | post | US share (pp) | F | -3.12 (0.61) | 39,436 |
| High nih dependence | post - pre | US share (pp) | R01 | 2.06 (1.65) | 45,066 |
| High nih dependence | post - pre | US share (pp) | F | -1.68 (2.81) | 45,066 |
| Low nih dependence | pre | US share (pp) | R01 | -3.55 (3.83) | 2,351 |
| Low nih dependence | pre | US share (pp) | F | 1.67 (2.84) | 2,351 |
| Low nih dependence | post | US share (pp) | R01 | 1.26 (1.04) | 13,076 |
| Low nih dependence | post | US share (pp) | F | -0.74 (1.05) | 13,076 |
| Low nih dependence | post - pre | US share (pp) | R01 | 4.64 (3.89) | 15,427 |
| Low nih dependence | post - pre | US share (pp) | F | -2.81 (2.69) | 15,427 |

**Data S1. (separate file)** The numbers behind every figure panel, one sheet per panel, with 95% confidence intervals and two-sided *P* values; a README sheet gives the source file and the command that produces each sheet.

## References

[1] R. K. Merton, Priorities in scientific discovery: A chapter in the sociology of science. American Sociological Review 22, 635–659 (1957).

[2] P. Dasgupta, P. A. David, Toward a new economics of science. Research Policy 23, 487–521 (1994).

[3] P. Stephan, How Economics Shapes Science (Harvard University Press, Cambridge, MA, 2012).

[4] R. J. Abdill, R. Blekhman, Tracking the popularity and outcomes of all bioRxiv preprints. eLife 8, e45133 (2019).

[5] N. Fraser, F. Momeni, P. Mayr, I. Peters, The relationship between bioRxiv preprints, citations and altmetrics. Quantitative Science Studies 1, 618–638 (2020).

[6] National Institutes of Health, Simplified Review Framework for NIH Research Project Grant Applications (2023). Notice NOT-OD-24-010, 19 October 2023; due dates on or after 25 January 2025.

[7] National Institutes of Health, Reporting Preprints and Other Interim Research Products (2017). Notice NOT-OD-17-050, 24 March 2017.

[8] openRxiv, Understanding DOIs for Preprints, https://openrxiv.org/dois-for-preprints/ (2025). Accessed 7 October 2026.

[9] J. Aschoff, Exogenous and endogenous components in circadian rhythms. Cold Spring Harbor Symposia on Quantitative Biology 25, 11–28 (1960).

[10] D. Ancona, C.-L. Chong, Entrainment: Pace, cycle, and rhythm in organizational behavior. Research in Organizational Behavior 18, 251–284 (1996).

[11] W. J. Orlikowski, J. Yates, It’s about time: Temporal structuring in organizations. Organization Science 13, 684–700 (2002).

[12] W. N. Espeland, M. Sauder, Rankings and reactivity: How public measures recreate social worlds. American Journal of Sociology 113, 1–40 (2007).

[13] medRxiv, Screening Procedures on medRxiv, https://connect.medrxiv.org/news/2022/06/13/screening_procedures (2022). 13 June 2022; accessed 9 October 2026.

[14] bioRxiv, Frequently Asked Questions, https://www.biorxiv.org/about/FAQ (2026). Accessed 9 October 2026.

[15] National Institutes of Health, NIH Data Book: Success Rates, R01-Equivalent and Research Project Grants, https://report.nih.gov/nihdatabook/category/10 (2025). Accessed 7 October 2026.

[16] N. Millar, B. Batalo, B. Budgell, Trends in the use of promotional language (hype) in abstracts of successful National Institutes of Health grant applications, 1985–2020. JAMA Network Open 5, e2228676 (2022).

[17] N. Millar, B. Batalo, B. Budgell, Promotional language (hype) in abstracts of publications of National Institutes of Health-funded research, 1985-2020. JAMA Network Open 6, e2348706 (2023).

[18] B. A. Jacob, L. Lefgren, The impact of research grant funding on scientific productivity. Journal of Public Economics 95, 1168–1177 (2011).

[19] C. J. G. Gersick, Time and transition in work teams: Toward a new model of group development. Academy of Management Journal 31, 9–41 (1988).

[20] A. E. Roth, J. K. Murnighan, F. Schoumaker, The deadline effect in bargaining: Some experimental evidence. American Economic Review 78, 806–823 (1988).

[21] R. Sever, M. Eisen, J. Inglis, Plan U: Universal access to scientific and medical research via funder preprint mandates. PLOS Biology 17, e3000273 (2019).

[22] Gates Foundation, Open Access Policy, Gates Foundation (2025). Effective 1 January 2025; accessed 30 September 2026.

[23] E. Zerubavel, Hidden Rhythms: Schedules and Calendars in Social Life (University of Chicago Press, Chicago, 1981).

[24] B. Latour, S. Woolgar, Laboratory Life: The Social Construction of Scientific Facts (Sage, Beverly Hills, 1979).

[25] National Institutes of Health, Revising the NIH/AHRQ/NIOSH Post-Submission Material Policy (2023). Notice NOT-OD-23-106, 31 March 2023; for the 25 May 2023 receipt date and beyond.

[26] National Institutes of Health, Update to the NIH/AHRQ/NIOSH Policy on Post-Submission Materials (2019). Notice NOT-OD-19-083, 15 February 2019.

[27] National Institutes of Health, NIH Removing AIDS Application Due Dates from NOFOs (2026). Notice NOT-OD-26-029, 28 January 2026; effective for January 2027 and later council rounds.

[28] National Institutes of Health, Update of NIH Late Application Submission Policy and End of Continuous Submission (2026). Notice NOT-OD-26-064, 31 March 2026; continuous submission applications accepted through 10 August 2026.

[29] J. Priem, H. Piwowar, R. Orr, OpenAlex: A fully-open index of scholarly works, authors, venues, institutions, and concepts. arXiv:2205.01833 (2022).

[30] National Institutes of Health, Submission Policies, https://grants.nih.gov/grants-process/submit/submission-policies (2026). Due dates that fall on a weekend, federal holiday or office closure move to the next business day; accessed 7 October 2026.

[31] National Institutes of Health, UPDATE: NIH Late Application Policy Due to Public Health Emergency for United States for 2019 Novel Coronavirus (COVID-19) (2020). Notice NOT-OD-20-091, 26 March 2020.

[32] National Institutes of Health, Updated Guidance on Reopening of NIH Extramural Activities Following the October 1, 2025 – Lapse in Appropriations (2025). Notice NOT-OD-26-012, 24 November 2025.

[33] National Institutes of Health, Standard Due Dates, https://grants.nih.gov/grants-process/submit/submission-policies/standard-due-dates (2026). Accessed 7 October 2026.

[34] C. A. Stewart, et al., Indiana University Pervasive Technology Institute, Indiana University, Bloomington, IN (2017).

